# Temperature Specific Regulation of NDR Kinase Orb6 by MAP kinase Sty1 to Promote Heat Stress Resilience

**DOI:** 10.1101/2021.02.26.432566

**Authors:** Laura P. Doyle, Robert N. Tams, Chuan Chen, Illyce Nuñez, Patrick Roman Haller, Fulvia Verde

## Abstract

Cellular response to environmental fluctuations, such as increased temperature, is crucial in promoting cell survival and plays an increasingly recognized role in cancer biology. Important cellular functions altered by heat stress are cell polarization and protein translation. Previous studies have shown that heat stress alters the dynamics of Cdc42, a key regulator of cell polarization in eukaryotes, and promotes RNP granule formation, reprogramming protein translation. The biological mechanisms underlying these vast changes are only partially known. Here, we report that conserved NDR kinase Orb6, a homologue of mammalian STK38, responds to heat stress and regulates heat stress resilience by modulating Cdc42 dynamics and promoting stress granule assembly. Also, we discovered a finely tuned mechanism whereby stress-activated MAP kinase Sty1 negatively regulates Orb6 kinase and Orb6 C-terminal phosphorylation during heat stress. Orb6 inhibition by Sty1 increases the sensitivity of the cell to heat stress in a temperature-specific manner, fostering increased stress resilience and metabolic adaptation. These observations highlight the role of NDR kinase in the process of heat adaptation and thermotolerance during environmental cell exposure to elevated temperatures.

**Summary statement:** Nuclear Dbf2-related kinase Orb6 inhibition by stress-activated protein kinase Sty1 promotes heat stress resilience in a temperature specific manner.

## Introduction

The ability to respond to varying environmental conditions, such as elevated temperature, is a vital cellular defense mechanism that promotes adaptation and survival (Zayani *et al*., 2025). Heat stress response activates adaptive mechanisms including antioxidant defenses, metabolic shifts, and heat shock proteins (HSPs) induction (Jung *et al*., 2013; Gomez-Pastor, Burchfiel and Thiele, 2018; Ciocca, Arrigo and Calderwood, 2013; Hu *et al*., 2022; Ding and Gao, 2025). Activation of the heat shock response usually provides cytoprotective effects, and HSPs have been found to promote survival of cancer cells (Wu *et al*., 2017). An increase in HSP levels has also been found in chronic inflammatory conditions such as rheumatoid arthritis (Fouani *et al*., 2022). In eukaryotes, severe heat stress (typically ≥ 42°C) rapidly disrupts cellular function, triggering a robust heat shock response, protein denaturation and aggregation, as well as a decrease in protein synthesis.

During heat stress, cells undergo distinct morphological changes. Mammalian cells often lose their spread morphology and become more rounded due to cytoskeletal reorganization (Velichko *et al*., 2013; Dressler *et al*., 2005; Gungor *et al*., 2014). However, the mechanisms that regulate morphological changes during heat stress are poorly understood. In the fission yeast *Schizosaccharomyces pombe*, an amenable model system, mild, sublethal heat stress at 36°C alters cell polarization and the dynamic behavior of Cdc42 GTPase (Vjestica *et al*., 2013), a key regulator of cell morphology in eukaryotes (Schwamborn and Püschel, 2004; Stankiewicz and Linseman, 2014). In unstressed conditions, Cdc42 displays oscillatory dynamics at cell tips (Das *et al*., 2012). During heat stress (Vjestica *et al*., 2013) as well as in response to different environmental conditions such as pheromone exposure (Bendezú and Martin, 2013), oxidative stress (Salat-Canela *et al*., 2021), or nitrogen starvation (Chen *et al*., 2019), an alternative “exploratory” pattern of Cdc42 emerges, where dynamic patches of active Cdc42 appear and disappear throughout the cell membrane. While the emergence of exploratory Cdc42 dynamic involves the activation of the conserved stress-activated MAP kinase Sty1/Spc1 (Salat-Canela *et al*., 2021; Mutavchiev, Leda and Sawin, 2016; Doyle *et al*., 2025), the mechanism that regulates Cdc42 dynamics shift during thermal stress is insofar poorly understood.

During heat stress, cells also rapidly suppress global protein synthesis to conserve energy and prevent accumulation of misfolded protein by the inhibition of translation initiation and mRNA sequestration into stress granules and P-bodies for storage or decay (Kedersha *et al*., 2005; Grousl *et al*., 2009). Stress granules (SGs) are membraneless cytoplasmic aggregates that form in response to various stress conditions, including heat shock, oxidative stress, osmotic stress, and infections (Marcelo *et al*., 2021). Their primary role is to protect mRNA and regulate translation during stress, helping cells survive adverse conditions. The formation process relies on liquid–liquid phase separation (LLPS), driven by multivalent interactions between RNA-binding proteins (RBPs) and untranslated mRNAs. Stress granules are critical adaptive structures that integrate translational control with cellular stress signaling (Desai *et al*., 2026). Dysregulation of SG dynamics is increasingly recognized as a contributor to human disease, and is strongly linked to neurodegeneration (ALS, FTD), cancer and viral infections (Ramaswami, Taylor and Parker, 2013; Anderson, Kedersha and Ivanov, 2015; Aguzzi and Altmeyer, 2016; Gilks *et al*., 2004; Alberti and Dormann, 2019).

Nuclear Dbf2-related (NDR) kinases are highly conserved members of the AGC protein kinase family which regulate essential cellular processes including morphogenesis, growth and proliferation, mitosis, and apoptosis (Hergovich *et al*., 2006; Cornils *et al*., 2011; Yang *et al*., 2014; Vichalkovski *et al*., 2008; Verde, Mata and Nurse, 1995; Chiba *et al*., 2009; Hergovich *et al*., 2007; Hergovich *et al*., 2009; Verde, Wiley and Nurse, 1998). Several studies have implicated dysregulation of NDR kinases in the development of cancer, and NDR also plays a role in neuronal differentiation (Adeyinka *et al*., 2002; Millward *et al*., 1998; Hauschild *et al*., 1999; Ross *et al*., 2000; Cornils *et al*., 2010; Zhang *et al*., 2015; Napoletano *et al*., 2011). The fission yeast *S. pombe* has a NDR/LATS kinase known as Orb6, which is responsible for regulation of polarized cell growth (Verde, Mata and Nurse, 1995). We previously reported that Orb6 kinase has separate roles in the control of Cdc42 GTPase (Das *et al*., 2015; Das *et al*., 2009), and mRNA translation (Nunez *et al*., 2016; Chen *et al*., 2019). Orb6 governs cell polarity by promoting the Ras1-dependent Cdc42 control axis (Chen *et al*., 2019) and by inhibiting a stress-activated Cdc42 module (Doyle *et al*., 2025; Salat-Canela *et al*., 2021) composed of the guanine nucleotide exchange factor Gef1 (Das *et al*., 2015; Das *et al*., 2012; Das *et al*., 2009; Coll *et al*., 2003) and the Cdc42 GAP (GTPase Activating Protein) Rga3 (Doyle *et al*., 2025; Gallo Castro and Martin, 2018). Downregulation of Orb6 kinase, which suppresses Ras1 GTPase and derepresses the stress-activated Cdc42 module, leads to the emergence of Cdc42 exploratory dynamics (Chen *et al*., 2019; Das *et al*., 2009).

Orb6 kinase also promotes polarized cell growth by spatiotemporal regulation of RNA-binding protein Sts5 (Nunez *et al*., 2016; Chen *et al*., 2019). Sts5 is an RNA binding protein (RBP) homologous to human Dis3L2, which is associated with Perlman’s syndrome and Wilm’s tumor in humans (Astuti *et al*., 2012; Lv *et al*., 2015; Robinson *et al*., 2015; Malecki *et al*., 2013; Toda *et al*., 1996; Vaggi *et al*., 2012; Jansen *et al*., 2009; Kurischko *et al*., 2011). Sts5 functions to repress translation of specific mRNAs, many of which encode proteins involved in polarized cell growth, via binding and sequestration into ribonucleoprotein (RNP) granules (Nunez *et al*., 2016), under conditions of nutritional stress (Nunez *et al*., 2016; Chen *et al*., 2019; Magliozzi and Moseley, 2021). Environmental stress such as nitrogen starvation, or crowded growth conditions leading to stationary phase, inhibit Orb6 kinase activity, which results in the translational repression of mRNAs involved in polarized growth via formation of Sts5 granules (Nunez *et al*., 2016; Chen *et al*., 2019).

In this paper, we report that heat stress downregulates Orb6 kinase activity, promoting Cdc42 exploratory dynamics and Sts5 coalescence into stress granules. Further, we find that Orb6 kinase inhibition increases stress granule formation and promotes cell resilience to heat stress. Recently, we reported that stress activated MAP kinase Sty1 negatively regulates Orb6 kinase activity during nitrogen starvation (Doyle *et al*., 2025). In this paper, we show that MAP kinase Sty1 has also a role in the induction of exploratory Cdc42 dynamics, Sts5 granule assembly, and the inhibition of Orb6 kinase activity during mild heat stress. Furthermore, we find that the levels of phosphorylation of Orb6-Thr456, the C-terminal hydrophobic motif, are decreased upon heat stress and that Sty1 activation enhances the sensitivity of Thr456 phosphorylation and Orb6 kinase function to heat stress. Thus, our observations identify a temperature specific regulation of NDR kinase Orb6 by MAP kinase Sty1 to promote exploratory Cdc42 dynamics and stress resilience.

## Results

### Orb6 kinase function and activity are downregulated by heat stress

We have previously shown that Orb6 kinase is sensitive to nutritional stress, and that downregulation of Orb6 kinase activity alters cell morphology and protein translation during nitrogen deprivation (Chen *et al*., 2019). During nutritional stress, Orb6 kinase activity decreases and therefore phosphorylation of the Cdc42 control factors and Orb6 substrates Gef1 (Cdc42 GEF) and Rga3 (Cdc42 GAP) declines, promoting exploratory Cdc42 dynamics (Das *et al*., 2015; Chen *et al*., 2019; Doyle *et al*., 2025). The onset of exploratory Cdc42 dynamics and increased Gef1 membrane localization were previously reported to occur also during exposure to sublethal mild heat stress at 36°C (Vjestica *et al*., 2013). Therefore, we investigated if Orb6 kinase has a role in the cellular response to heat stress. Consistent with previous findings, we found that Gef1-3YFP localizes to the cell membrane after 30 minutes at 36°C (Fig. 1A, B) as compared to the control which remained at 25°C. Further, we found that Gef1-3YFP also localizes to the cell membrane at the higher, severe heat stress temperature of 42°C (Fig. 1A, B).

**Figure 1:**
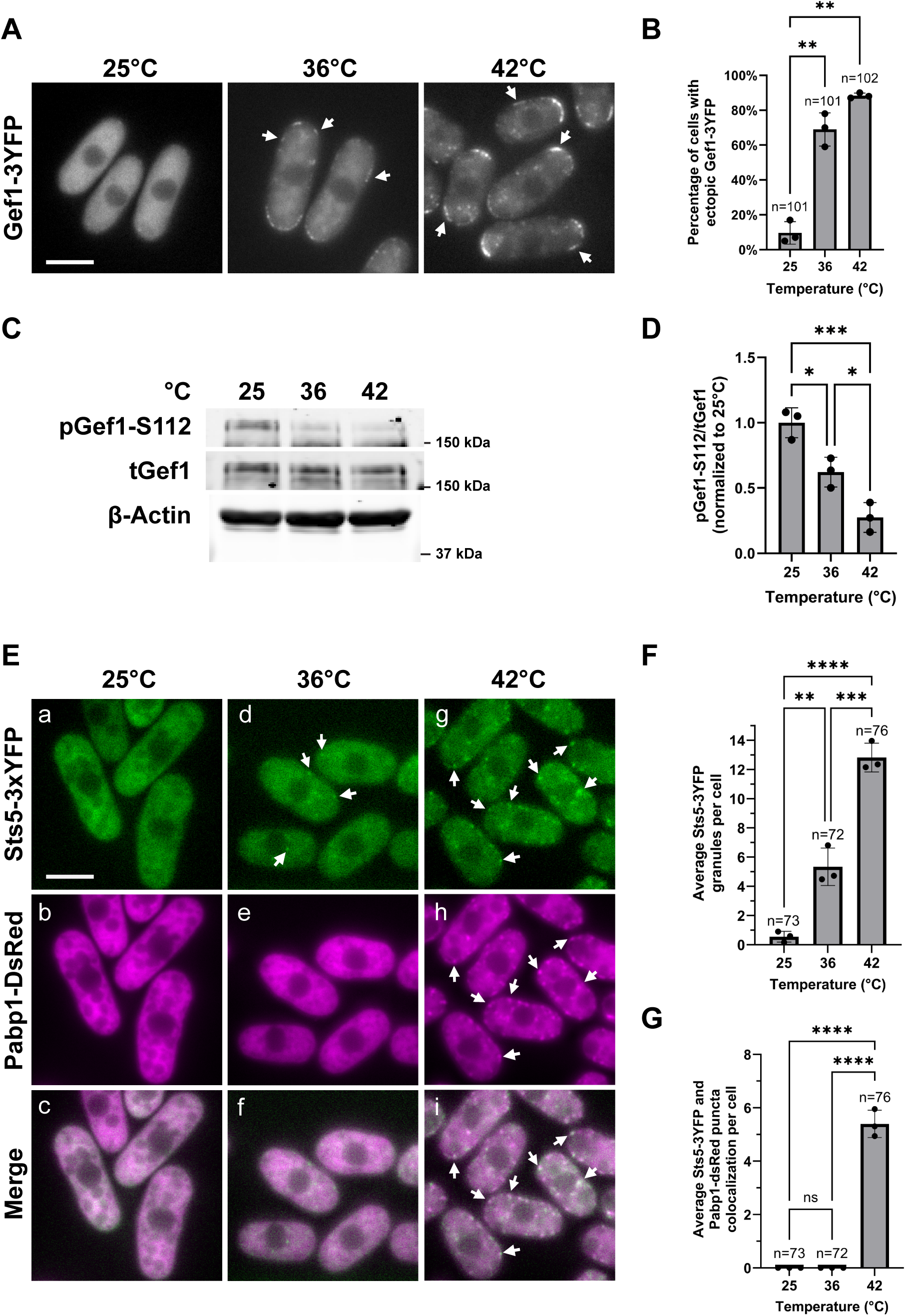
Orb6 activity is negatively regulated by heat stress. (A) Gef1-3YFP localizes the cell membrane upon heat exposure at 36°C or 42°C for 30 minutes (arrows). Scale bar, 5 μm. (B) Quantification of percentage of cells that display ectopic Gef1-3YFP localization from A based on three independent experiments. Data presented as mean ± SD, *p* values are determined by one-way ANOVA with Tukey’s HSD test p ≤ 0.01, **. n = number of cells quantified. (C) Orb6 activity, as measured by Gef1-S112 phosphorylation, decreases after 30 minutes at 36°C or 42°C. (D) Quantification of pGef1-S112/tGef1 from C upon temperature stress exposure based on three independent experiments. Data are presented as in B, p ≤ 0.05, *; p ≤ 0.001, ***. (E) *Sts5-3xGFP pabp1-DsRed* cells were exposed to 36°C or 42°C for 30 minutes. At 36°C, Sts5-3xGFP formed cytoplasmic puncta (d, arrows) but no stress granule formation was visible (f). At 42°C, Sts5 partially co-localized with the stress granule marker Pabp1-DsRed (g-i, arrows). Scale bar, 5 μm. (F) Quantification of average number of Sts5 puncta present in each cell from E based on three independent experiments. Data are presented as in B, p ≤ 0.01, **; p ≤ 0.001, ***, p ≤ 0.0001, ****. n = number of cells quantified. (G) Quantification of average number of Sts5 puncta that colocalized by stress granule marker Pabp1 in each cell from E based on three independent experiments. Data are presented as in B, p ≤ 0.0001, ****. n = number of cells quantified.

As Gef1 is a substrate that is phosphorylated by Orb6 at serine 112, we sought to measure Orb6 activity using a previously described custom phospho-specific antibody for Gef1-S112 (Chen *et al*., 2019). We found that Gef1-S112 phosphorylation decreases after 30 minutes at 36°C, a sublethal temperature that induces heat stress, and is further exacerbated at 42°C, a lethal temperature that induces a severe heat shock response (Fig. 1C,D). These results indicate the Orb6 activity decreases during both mild and severe heat stress exposure, fostering exploratory Cdc42 dynamics.

During heat exposure, cells reorganize their RNA and protein metabolism by forming two major types of cytoplasmic granules: Stress Granules (SGs) (Guzikowski, Chen and Zid, 2019; Mahboubi and Stochaj, 2017), and Processing Bodies (P-bodies). Both are membraneless organelles formed by liquid–liquid phase separation, but they have distinct roles and compositions. We previously demonstrated that Orb6 kinase controls polarized cell growth and protein translation by phosphorylating the highly conserved RNA binding protein Sts5. Sts5, an Orb6 substrate, remains diffused in the cytoplasm when Orb6 is active, whereas it partially co-localizes with P-(Processing) Bodies following Orb6 inhibition, or in media lacking glucose or nitrogen (Nunez *et al*., 2016; Magliozzi and Moseley, 2021).

Consistent with Orb6 kinase activity being downregulated by heat exposure, we found that Sts5-3YFP remains diffused in the cytoplasm at 25°C, whereas it forms puncta at both 36°C and 42°C (Fig. 1E, F). We found that Sts5-3YFP puncta readily associate with stress granules (using Pabp1-dsRed stress granule marker) at 42°C (Figure 1E,g-i, 1G), but not at 36°C where we find no stress granule assembly (Figure 1E,d-f, 1G). Conversely, we find that Sts5-3YFP colocalizes with P-body marker Dcp1-mCherry at both 36°C and 42°C (Fig. S1).

We sought to further confirm Sts5 and stress granule co-localization, under conditions of glucose limitation, which strongly induces stress granules formation (Nilsson and Sunnerhagen, 2011) as well as Sts5 aggregation into RNP granules (Nunez *et al*., 2016). We investigated the localization of *ssp1* mRNA, a Sts5-bound mRNA which is translationally repressed upon formation of Sts5 puncta (Nunez *et al*., 2016) by RNA-FISH analysis. *ssp1* encodes a CamK kinase (Hanyu *et al*., 2009; Matsusaka *et al*., 1995).

We observed co-localization of Sts5 puncta, *ssp1* mRNA, and stress granules in cells cultured in medium lacking glucose (Fig. 2A). We previously used *ssp1* mRNA to detect co-localization of Sts5 with P-bodies (Nuñez *et al*., 2016).

**Figure 2:**
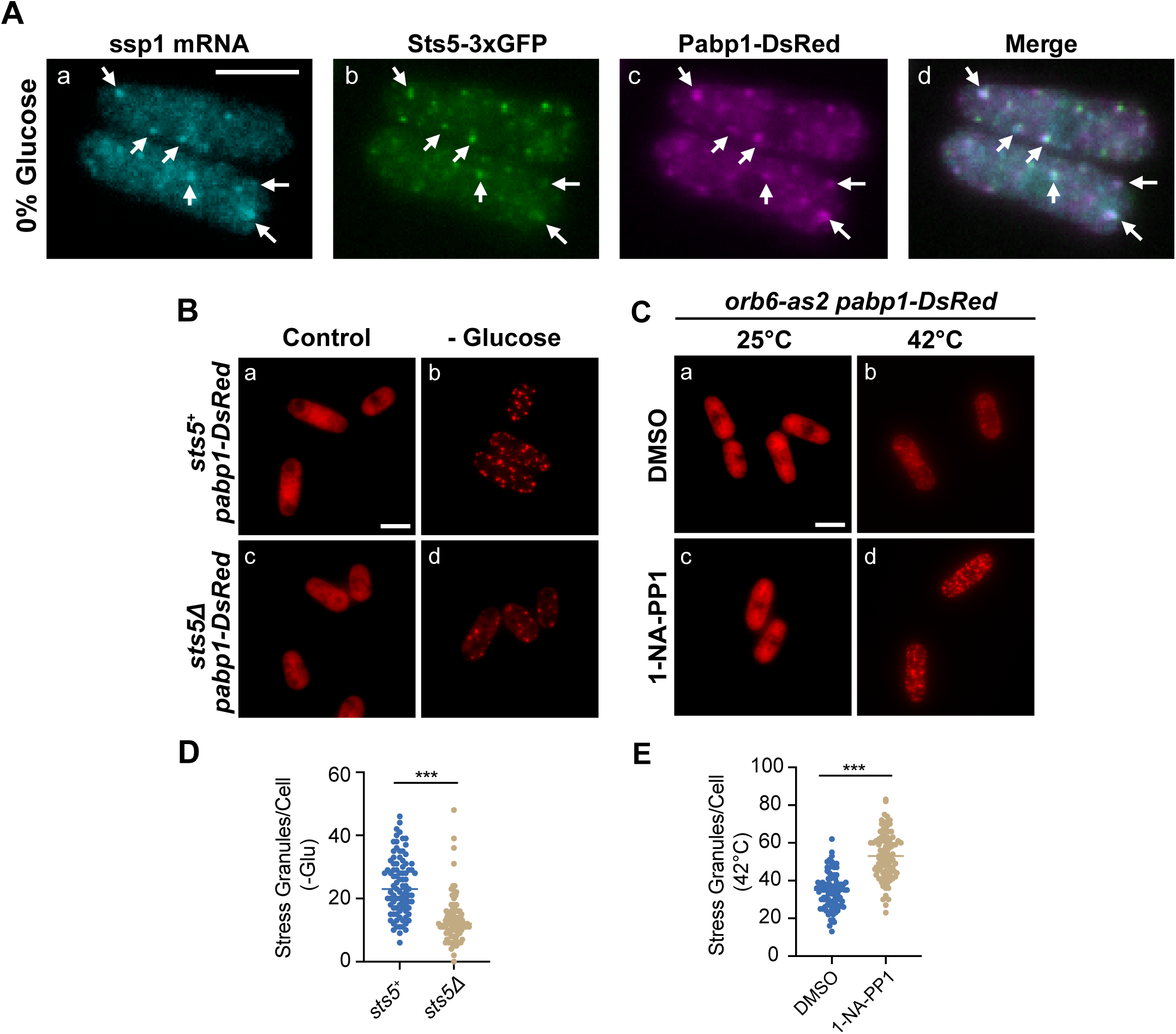
Orb6 and Sts5 regulate stress granule formation. (A) RNA FISH visualization of ssp1 mRNA in fixed cells cultured for 20 minutes in minimal medium containing 0% glucose. Hybridization of RNA was performed with 20-mer DNA oligonucleotides (Stellaris) labeled with Quasar 705 fluorochromes. ssp1 mRNA co-localized with Sts5-3xGFP and the stress granule marker Pabp1-DsRed (arrows). Images are deconvolved projections from Z-stacks (12 slices) separated by a step size of 0.3 μM (Bar = 5 μM). (B) *pabp1-DsRed* and *sts5Δ pabp1-DsRed* cells were cultured in EMM medium with (a, c) or without (b, d) glucose for 20 minutes. In untreated controls, the stress granule marker Pabp1-DsRed remained diffuse throughout the cytoplasm. Following glucose deprivation, *sts5Δ* cells formed less stress granules than the control (b, d). Representative images are deconvolved projections from Z-stacks separated by a step size of 0.3 μM (Bar = 5 μM). (C) Inhibition of Orb6 primes cells for the formation of stress granules. *orb6-as2 pabp1-DsRed* cells were cultured in minimal medium containing adenine, and 15 minutes prior to heat shock 1-NA-PP1 inhibitor was added, or DMSO for controls. In cells maintained at 25°C, Pabp1-DsRed remained diffuse within the cytoplasm (a, c). However, in cells that were heat shocked for 20 minutes at 42°C (15 minutes post addition of 1-NA-PP1), there was an increase in stress granule formation compared to the DMSO control (b, d). Representative images are deconvolved projections from Z-stacks separated by a step size of 0.3 μM (Bar = 5 μM). (D) The number of stress granules were quantified for experiment represented in B (n=90 cells/strain/condition total, performed in biological triplicate), and there was a statistically significant decrease in stress granule formation for the *sts5Δ* mutant as compared to the control (***p<0.001; unpaired t-test). (E) The increase in stress granule formation from C was statistically significant and performed in biological triplicates (n=90 cells/strain/condition total; ***p<0.001, unpaired t-test).

Overall, these findings show that Orb6 kinase activity is downregulated upon heat stress and heat shock, fostering Cdc42 exploratory dynamics and coalescence of RNA binding protein Sts5 into stress granules and P bodies.

### Orb6 inhibition and Sts5 Aggregation Promote Stress Granule Assembly

We have previously demonstrated that Orb6 inhibition promotes the formation of P-Bodies in a Sts5 dependent manner (Nunez *et al*., 2016), suggesting that Sts5 aggregates may promote P-Body formation during glucose starvation. To test whether Sts5 plays a role in stress granule nucleation, we followed the formation of stress granules in the *sts5Δ* mutant using the stress granule marker Pabp1-DsRed. In this particular experiment, we induced stress granules via glucose deprivation, which strongly induces stress granules assembly, facilitating quantification. We found that the number of stress granules decreased by 44% in *sts5Δ* mutant cells as compared to control cells (Fig. 2B, b, d; Fig. 2D) cultured for 20 minutes in media lacking glucose. In media containing glucose, Pabp1-DsRed is largely diffuse within the cytoplasm in both the *sts5Δ* and control strains (Fig. 2B; a, c).

To determine if inhibition of Orb6 kinase drives stress granule formation, we treated *orb6-as2 pabp1-DsRed* cells with 1-NA-PP1 inhibitor and found that inhibition of Orb6-as2 at 25°C does not induce stress granule formation (Fig. 2C; a, c), These findings indicate that Orb6 kinase inhibition alone is not sufficient to induce stress granule formation, while it is sufficient to promote P-Body formation, as we previously reported (Nunez *et al*., 2016). However, when 1-NA-PP1 treated cells were subsequently heat shocked for 20 minutes at 42°C, we observed a remarkable increase in the number of stress granules, as compared to the heat shocked DMSO control (Fig. 2C; b, d). The number of stress granules formed was quantified for three independent experiments (n=90 cells/condition), and the number of stress granules increased, on average, by 54% for the 1-NA-PP1 treated cells as compared to the heat shocked DMSO control (Fig. 2E). Overall, these data indicate that the presence of Sts5 is important for the formation of stress granules, and that Orb6 kinase plays a role in this process by modulating the extent of stress granule formation.

### Sts5 aggregation Modulates Heat Resilience in *S. pombe*

Stress granule formation provide a mechanism to protect cells from environmental stress and promote cell survival (Mahboubi and Stochaj, 2017). These condensates can modulate susceptibility to heat stress (Riback *et al*., 2017; Kroschwald and Alberti, 2017). Since Sts5 responds to heat by co-localizing with both stress granules and P-Bodies (Fig. 1E, Fig. S1) and is important for stress granule formation (Fig. 2B), we hypothesized that modulating the assembly of Sts5 into RNP granules plays a role in cell protection from heat stress. Sts5 contains an intrinsically disordered domain (IDD), which is phosphorylated by Orb6 kinase at serine 86 (Chen *et al*., 2019). Modification of Sts5 serine 86 to alanine promotes increased Sts5 puncta formation, increases cell resilience during nutritional stress and promotes chronological lifespan extension (Chen *et al*., 2019). Thus, we tested if loss of Sts5 (*sts5Δ*), or the presence of the *sts5-S86A* mutation affect cell survival during extended heat stress or heat shock. To test the role of Sts5 during heat stress and in heat tolerance (exposure to heat for extended times), wild-type, *sts5Δ*, *sts5-HA*, and *sts5-S86A-HA* log-phase cells were incubated for 6 hours at heat stress temperature (36.5°C) or control temperature (25°C) in a shaking incubator and plated to calculate colony forming units (CFUs) (see Methods). We found that the *sts5Δ* strain exhibited decreased survival compared to wild-type (Fig. 3A). Conversely, the *sts5-S86A-HA* strain displayed increased survival, as compared to the *sts5-HA* or wild-type controls (Fig. 3A). The *sts5-HA* strain exhibited similar survival as compared to the wild type, as expected. We previously established that addition of the *sts5-S86A* mutation does not alter Sts5 protein levels (Chen *et al*., 2019).

**Figure 3:**
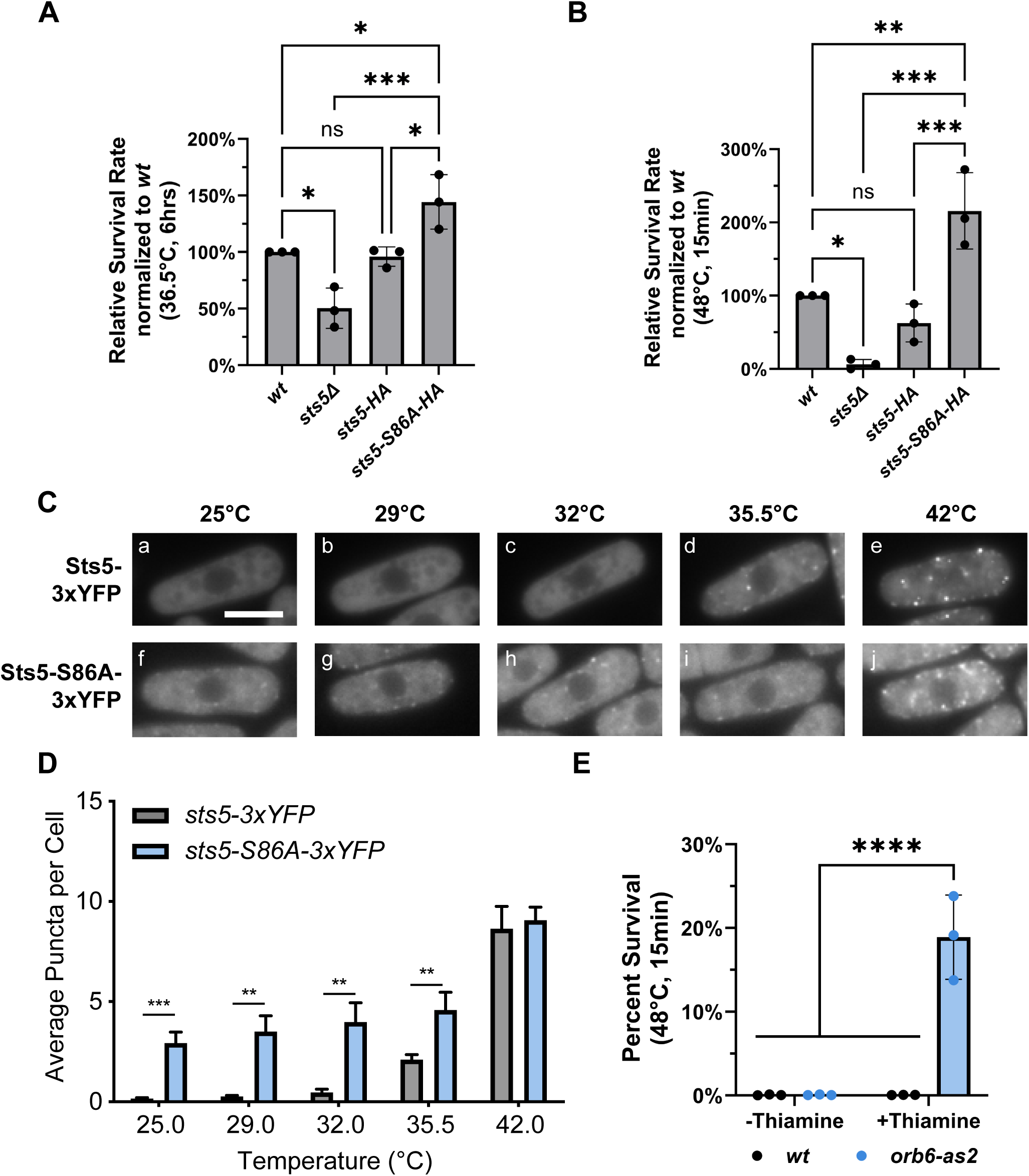
Orb6 and Sts5 modulate survival following heat stress. (A) *Wild-type, sts5Δ, sts5-HA,* and *sts5-S86A-HA* cells were grown for 6 hours at either 25°C or 36.5°C. Following prolonged heat stress at 36.5°C, s*ts5Δ* cells displayed decreased survival compared to wild type, whereas *sts5-S86A-HA* showed increased survival compared to the *sts5-HA* control. Data are presented as mean ± SD, p values determined by one-way ANOVA with Tukey’s HSD test p ≤ 0.05, *; p ≤ 0.01, **; p ≤ 0.001, ***. (B) The experiment in A was repeated, but cells were instead incubated for 15 minutes at 48°C temperature, while untreated controls were incubated at 25°C. Again, *sts5Δ* cells exhibit decreased survival, whereas the *sts5-S86A-HA* mutant exhibited increased survival. Data are presented as mean ± SD, p values determined by one-way ANOVA with Tukey’s HSD test p ≤ 0.05, *; p ≤ 0.001, ***. (C) In the *sts5-S86A-3xYFP* mutant, the number of Sts5 granules were increased at temperatures in which the *sts5-3xYFP* control did not form granules. (D) Quantification from results obtained in C. There is a statistically significant increase in average puncta per cell for the *sts5-S86A-3xYFP* strain as compared to the control for cells incubated between 25°C and 35.5°C Data are presented as mean ± SD, p values determined by two-way ANOVA with Tukey’s HSD test p ≤ 0.01, **; p ≤ 0.001, ***. n=60 cells/strain performed in biological triplicates (E) *Wild-type* and *orb6-as2* strains were cultured in the presence (+Thiamine) or absence of thiamine (-Thiamine), heat shocked, and plated. Inhibition of Orb6 results in a 17-fold increase in survival as compared to the untreated control. Data are presented as mean ± SD, p values determined by two-way ANOVA with Tukey’s HSD test p ≤ 0.0001, ****.

To test response upon acute heat shock, wild-type, *sts5Δ*, *sts5-HA*, and *sts5-S86A-HA* strains were exposed to a 48°C heat shock for 15 minutes. We found that the *sts5Δ* strain exhibited decreased survival as compared to the wild-type control (Fig. 3B). Again, the *sts5-S86A-HA* mutant exhibited increased survival as compared either the wild-type or *sts5-HA* controls (Fig. 3B).

Thus, our data indicate that Sts5 coalescence is important for survival and heat stress resilience following various durations and intensities of exposure to elevated temperatures. Therefore, we tested whether Sts5 granule assembly is modulated by increased temperature and whether the *sts5-S86A* mutation increases formation of Sts5 puncta following exposure to heat. To do this, *sts5-3xYFP* and *sts5-S86A-3xYFP* strains were cultured at 29°C, 32°C, 35.5°C, or 42°C for 30 minutes, while control *sts5-3xYFP* and *sts5-S86A-3xYFP* cells were maintained at 25°C throughout the experiment. Following heat stress, fluorescence microscopy was performed and the number of Sts5 puncta were counted at each temperature. The *sts5-3xYFP* control began to form visible puncta at 35.5°C (Fig. 3C; a-d), whereas the *sts5-S86A-3xYFP* mutant began to form puncta already at 25°C (Fig. 3C, f-j). These differences were statistically significant (Fig. 3D), and *sts5-S86A* predisposition to the formation of puncta was readily apparent until the temperature reached 42°C at which point the control formed approximately the same number of puncta (Fig. 3Ce, Cj, 3D).

Our findings indicate an important role for serine 86 phosphorylation in modulating the extent of Sts5 puncta assembly in response to increased temperatures (Fig. 3C, D). Since Orb6 kinase phosphorylates serine 86 (Chen *et al*., 2019), we tested if downregulation of Orb6 kinase also increases cell resilience to heat stress. To transcriptionally repress Orb6, a thiamine-repressible o*rb6-as2* strain was used. Log phase wild-type and *orb6-as2* cells were grown with or without 15 µM thiamine for 16 hours at 32°C. Samples were diluted to an equivalent optical density and were heat shocked at 48°C for 15 minutes while control samples were maintained at 32°C. The *orb6-as2* strain, in the presence of thiamine, exhibited a striking increase in survival following heat shock compared to the untreated control (Fig. 3E). This striking increase in resilience was readily observed on Petri plates (Fig. S2) and was notably higher than the relative increase in heat resilience observed for the *sts5-S86A* mutant (Fig. 3A-B). Overall, these data indicate that the Orb6 kinase modulates stress granule assembly and resilience to heat stress, in part, by phosphorylating mRNA binding protein Sts5 on serine 86.

### Conserved MAP kinase Sty1 negatively regulates Orb6 activity in a temperature specific manner

The mitogen-activated protein kinase (MAPK) Sty1/Spc1 is activated upon exposure to diverse stressors such as nitrogen starvation, osmotic changes, oxidative stress, and heat shock (Toone and Jones, 1998; Shiozaki and Russell, 1996; Millar, Buck and Wilkinson, 1995; Degols, Shiozaki and Russell, 1996). Recently, we have shown that Sty1 activation during nitrogen starvation negatively regulates Orb6 activity (Doyle *et al*., 2025). Therefore, we tested if Sty1 has a role in Orb6 regulation during heat stress, by visualizing Sts5-3YFP aggregation during either mild or severe heat stress (36°C or 42°C) for 30 minutes in the presence or absence of *sty1.* We find that upon loss of *sty1,* Sts5-3YFP does not assemble into granules at 36°C but it still assembles into granules at 42°C (Fig. 4A). The number of Sts5 granules per cell at 42°C is similar in both the presence or absence of *sty1* (Fig. 4B). These results indicate that Sts5 granule assembly during heat stress is dependent on the presence of Sty1 at 36°C but not at 42°C.

**Figure 4:**
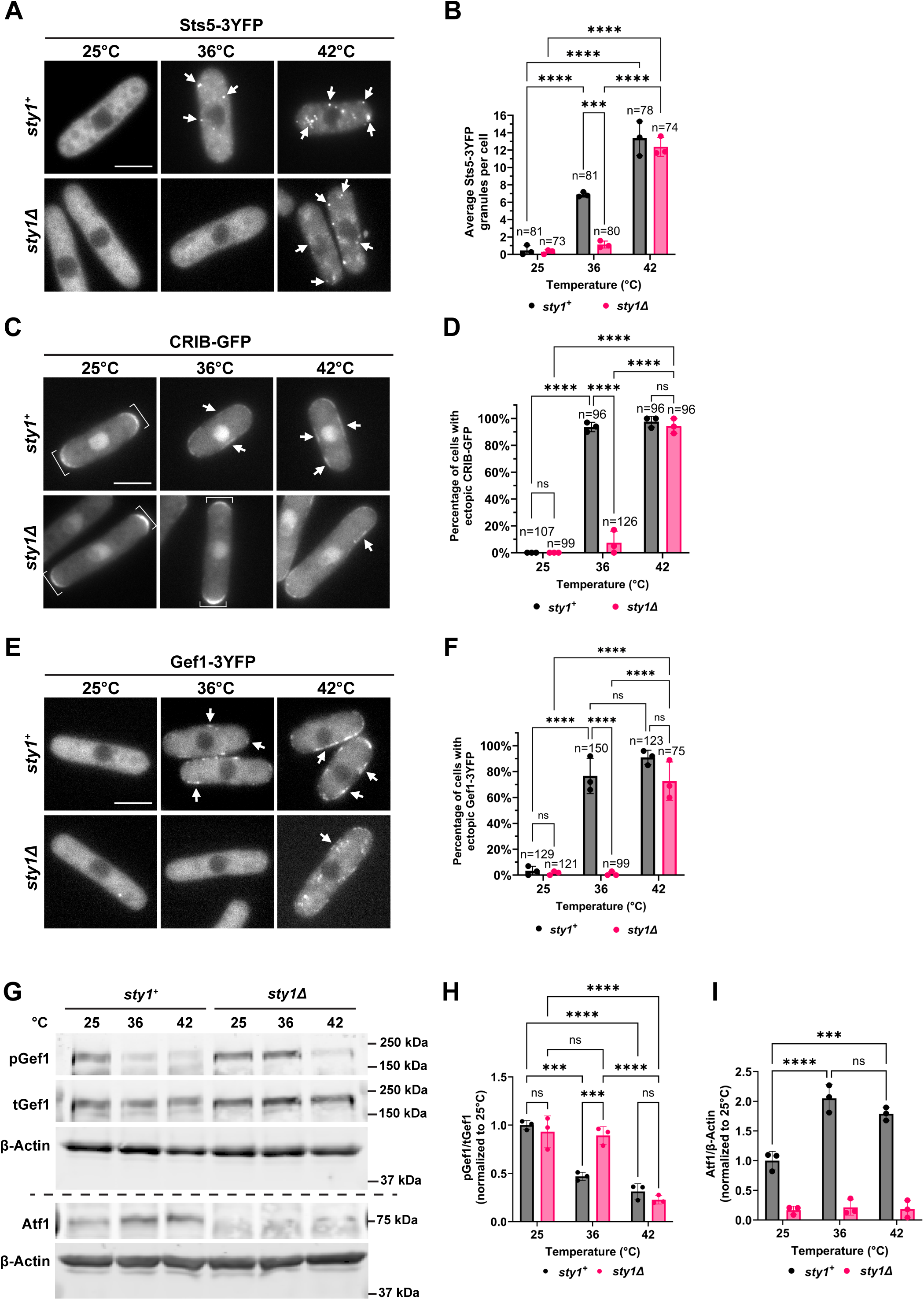
The distribution of Sts5, Gef1, and active Cdc42 is temperature sensitive and is modulated by Sty1 in a temperature specific manner. (A) Sts5-3YFP forms puncta after 30 minutes at 36°C and 42°C (arrows). In the absence of *sty1,* Sts5-3YFP only clusters at 42°C. Images are a max projection Z-stack of 6 images separated by a step-size of 0.3 μm. Scale bar, 5 μm. (B) Quantification of average number of Sts5 puncta present in each cell from A based on three independent experiments. Data are presented as mean ± SD, p values determined by two-way ANOVA with Tukey’s HSD test p ≤ 0.001, ***; p ≤ 0.0001, ****. n = number of cells quantified. (C) Active Cdc42 forms exploratory patches upon exposure to 36°C and 42°C when *sty1* is present but only forms patches at 42°C when *sty1* is absent (arrows). Active Cdc42 is localized at the cell tips in control cells at 25°C and in *sty1Δ* cells at 25°C and 36°C (brackets) Images are a max projection Z-stack of 6 images separated by a step-size of 0.3 μm. Scale bar, 5 μm. (D) Quantification of percentage of cells that display ectopic CRIB-GFP localization from C based on three independent experiments. Data presented as in D, p ≤ 0.0001, ****. n = number of cells quantified. (E) Gef1-3YFP localizes to the cell membrane at 36°C and 42°C (arrows). Loss of *sty1* leads to Gef1-3YFP localization to the cell membrane only at 42°C. Scale bar, 5 μm. (F) Quantification of percentage of cells that display ectopic Gef1-3YFP localization from B based on three independent experiments. Data presented as in D, p ≤ 0.0001, ****. n = number of cells quantified. (G) Gef1-S112 phosphorylation by Orb6 decreases after 30 minutes at 36°C or 42°C. In *sty1Δ* deletion cells, pGef1-S112 remains constant at 36°C but decreases at 42°C. Atf1 levels increase upon exposure to 36°C or 42°C for 30 minutes in control cells. β-Actin was used as a loading control. (H) Quantification of pGef1-S112/tGef1 from A upon temperature stress exposure in control or *sty1Δ* deletion mutant cells based on three independent experiments. Data are presented as mean ± SD, *p* values are determined by two-way ANOVA with Tukey’s HSD test p ≤ 0.001, ***; p ≤ 0.0001, ****. (I) Quantification of Atf1/Actin from A upon temperature stress in control or *sty1Δ* deletion mutant cells based on three independent experiments. Data are presented as mean ± SD, *p* values are determined by two-way ANOVA with Tukey’s HSD test p ≤ 0.001, ***; p ≤ 0.0001, ****.

Previously, we have shown that Orb6 phosphorylates two regulators of Cdc42, a GEF Gef1 (Das *et al*., 2015) and a GTPase activating protein (GAP) Rga3 (Doyle *et al*., 2025) to spatially regulate Cdc42 dynamics. When Orb6 is inhibited during nitrogen starvation, Gef1 and Rga3 move to the cell membrane and facilitate the onset of exploratory Cdc42 dynamics, where active Cdc42 forms dynamic patches along the cell membrane (Doyle *et al*., 2025). These exploratory Cdc42 dynamics are also found upon different stress exposure, including heat stress at 36°C (Vjestica *et al*., 2013). Thus, we sought to investigate if exploratory Cdc42 dynamics were dependent on Sty1 upon heat exposure. We visualized active Cdc42 using fluorescently tagged Cdc42-Rac Interactive Binding domain bioreporter CRIB-GFP or the Cdc42 GEF Gef1 using Gef1-3YFP. We find that exploratory Cdc42 dynamics and Gef1-3YFP membrane localization are induced by heat exposure at 36°C and 42°C and that these effects are dependent on Sty1 at 36°C, but not 42°C (Fig. 4C-F).

Further, we tested the effects of heat stress and heat shock on Gef1-S112 phosphorylation by Orb6 kinase, in the presence and absence of *sty1.* We find that Gef1-S112 phosphorylation decreases upon exposure to 36°C and 42°C (Fig. 4G,H). In the absence of *sty1,* we find that Gef1-S112 phosphorylation remains constant at 36°C but decreases to similar levels as wild-type at 42°C (Fig. 4G, H). Sty1 activation in these experimental conditions was validated by measuring levels of Atf1, a transcription factor phosphorylated by Sty1 upon Sty1 activation (Shiozaki, Shiozaki and Russell, 1998; Wilkinson *et al*., 1996; Sun *et al*., 2024). As expected, we find that Atf1 levels increase upon cell exposure to 36°C or 42°C. (Fig. 4G, I), confirming that Sty1 is activated under these experimental conditions.

Overall, these results indicate that Sty1 kinase activation during temperature stress exerts a temperature specific effect on Orb6 kinase activity; Sty1 activation is necessary to inhibit Orb6 activity and biological functions at 36°C while is not necessary at higher, heat shock temperatures (42°C).

### Phosphorylation of the C-terminal hydrophobic motif Orb6-T456 is temperature sensitive

To better study the molecular mechanisms of Orb6 kinase regulation, we developed a phospho-specific antibody that recognizes Orb6 phosphorylation by an upstream kinase at the C-terminal hydrophobic motif site, Threonine 456 (Fig. 5A). Phosphorylation of the conserved phosphorylation site, Thr456, is essential for Orb6 function (Liu and Young, 2012). We validated the specificity of the phospho-Orb6-T456 antibody (hereafter referred to as pOrb6-T456) by performing a dot blot with 4 different peptides of Orb6. The pOrb6-T456 antibody was specific only to phosphorylated Threonine 456 and not to the corresponding unphosphorylated peptide or a similar sequence at the C-terminal, Threonine 464 (Fig. 5B). Previous studies have identified interactions between Nak1 kinase and Orb6 kinase suggesting that Nak1 functions upstream of Orb6 (Liu and Young, 2012). However, direct phosphorylation of Orb6 by Nak1 has yet to be reported. To test if Orb6-T456 phosphorylation is Nak1-dependent, we exposed temperature-sensitive mutant of *nak1 (nak1-ts/orb3-167)* (Snell and Nurse, 1994; Leonhard and Nurse, 2005) to restrictive temperature (36°C) for 3 hours to downregulate Nak1 activity and measured pOrb6-T456 levels in HA-tagged Orb6 (Fig. 5C). We find that pOrb6-T456 strongly decreases at 36°C in the *nak1-ts* mutant (Fig. 5C, D). Further, we find that pOrb6-T456 also decreases to a lesser extent, at 36°C in the control *nak1+* cells. (Fig. 5C, D). These results indicate the Orb6 phosphorylation at the C-terminal hydrophobic motif T456 site is dependent on Nak1. Our observations also suggest that Orb6-T456 phosphorylation is affected by heat stress.

**Figure 5:**
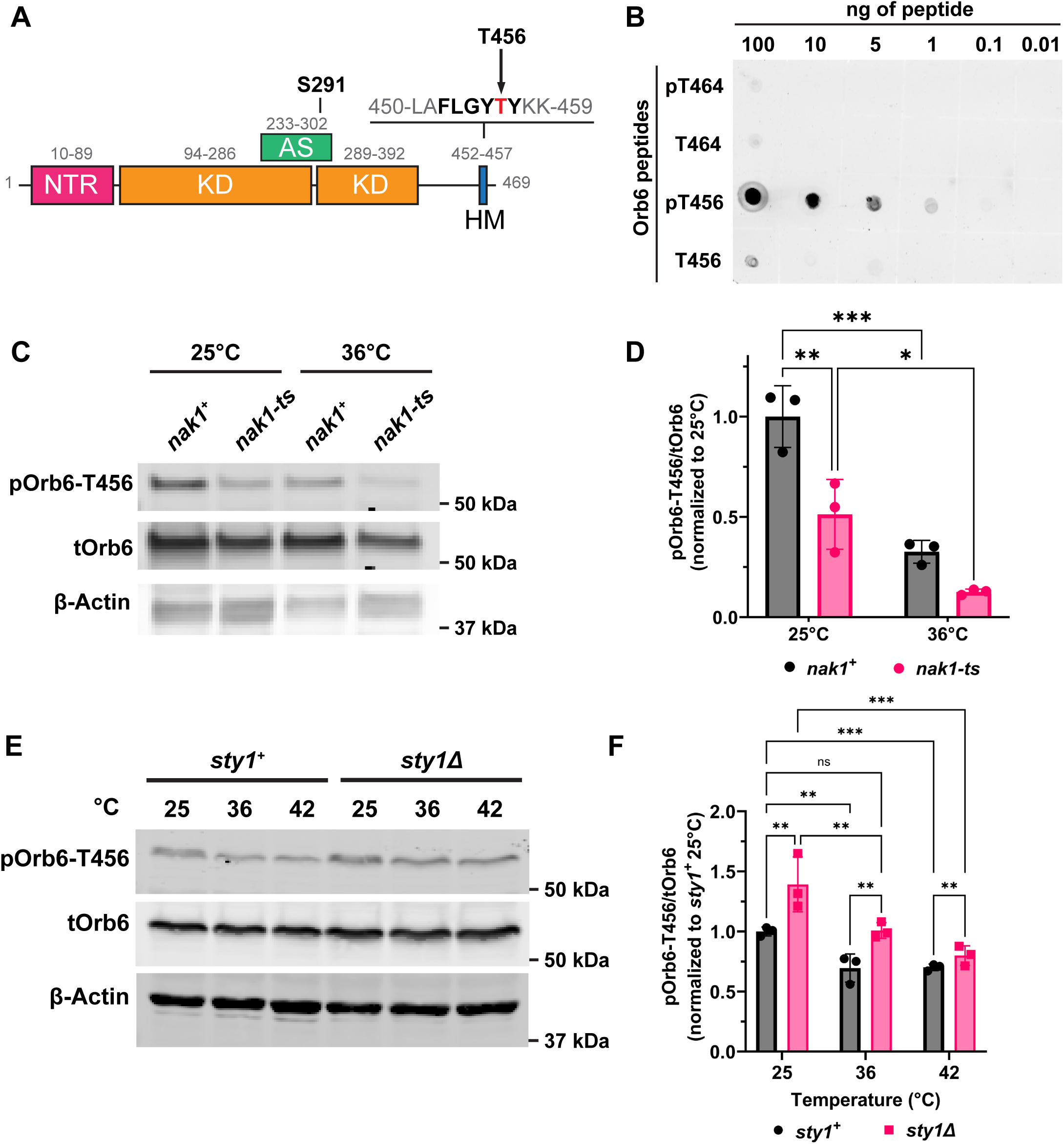
The phosphorylation state of the Orb6 C-terminal hydrophobic motif is temperature sensitive. (A) Protein domain map of Orb6 kinase highlighting two phosphosites: S291 in the activation segment and T456 in the hydrophobic motif. NTR:N-terminal region; KD: kinase domain; AS: activation segment; HM: hydrophobic motif. (B) Dot plot showing specificity of phosphor-Orb6-T456 antibody. (C) Orb6-456 phosphorylation decreases when temperature sensitive mutant of *nak1* (*nak1-ts)* is exposed to restrictive temperature for 3 hours. (D) Quantification of pOrb6-T456/tOrb6 from C based on three independent experiments. Data are presented as mean ± SD, *p* values are determined by two-way ANOVA with Tukey’s HSD test p ≤ 0.05, *; p ≤ 0.01, **; p ≤ 0.001, ***. (E) Orb6-T456 phosphorylation decreases after 30 minutes at 36°C or 42°C in control or *sty1Δ* deletion mutant cells. Orb6-T456 phosphorylation is higher in *sty1Δ* mutants. β-Actin was used as a loading control. (F) Quantification of pOrb6-T456/tOrb6 from A based on three independent experiments. Data are presented as mean ± SD, *p* values are determined by two-way ANOVA with Tukey’s HSD test p ≤ 0.01, **; p ≤ 0.001, ***.

To test if Sty1 kinase has a role in modulating Orb6-T456 phosphorylation, we exposed cells to 36°C or 42°C for 30 minutes in the presence or absence of *sty1.* We found that pOrb6-T456 phosphorylation decreases upon exposure to heat, in both control *sty1^+^* or *sty1Δ* deletion mutants (Fig. 5E-F). However, loss of *sty1* leads to overall increased levels of pOrb6-T456, as compared to control cells (Fig. 5E-F). This increase is particularly evident at 25°C and 36°C, indicating that Sty1 regulates the levels of phosphorylation of the Orb6 protein at the T456 site. Our observations suggest that the increased levels of Orb6 phosphorylation, in the absence of Sty1, play a role in maintaining sustained Orb6 kinase activity during heat stress at 36°C, thereby suppressing the onset of Cdc42 exploratory dynamics and the induction of Sts5 granule formation. This idea is supported by the observation that the extent of Gef1-S112 phosphorylation (a readout of Orb6 kinase activity) is similar in *sty1Δ* cells at 36°C and unstressed wild-type controls at 25°C. Since Orb6 T456 phosphorylation decreases in response to heat in both control and *sty1Δ* deletion mutant cells, our data also point to a Sty1-independent mechanism of downregulation of Orb6 activity in response to increasing heat stress.

## Discussion

### Conserved NDR kinase Orb6 activity is thermo-sensitive

Proper cellular response to environmental fluctuations, such as heat exposure, is critical for survival and is increasingly recognized as a key factor in human disease. Heat stress disrupts essential cellular processes, including cell polarization (Vjestica *et al*., 2013; Pawlik *et al*., 2013; Singh *et al*., 2016; Delley and Hall, 1999) and protein translation (Kedersha *et al*., 2005; Grousl *et al*., 2009). However, the underlying mechanisms mediating these effects are not completely understood.

Here, we demonstrate that the conserved fission yeast NDR kinase Orb6, homologous to mammalian STK38, is inactivated by heat stress, a process that alters the pattern of active Cdc42 distribution at the membrane and facilitates stress granule assembly. Overall, our findings provide experimental evidence explaining, at least in part, some of the physiological effects of heat stress. We also uncover a regulatory mechanism in which the stress-activated MAP kinase Sty1 negatively regulates Orb6 activity and Orb6 C-terminal phosphorylation during heat stress (Fig. 6). This inhibition sensitizes cells to temperature-specific heat stress, promoting adaptive metabolic responses and thermotolerance. These findings underscore the pivotal role of NDR kinase signaling in cellular adaptation to elevated temperatures. Further, our observations reveal a novel role for conserved stress activated MAP kinase Sty1 in RNP granule formation, mediated by Orb6 kinase.

**Figure 6:**
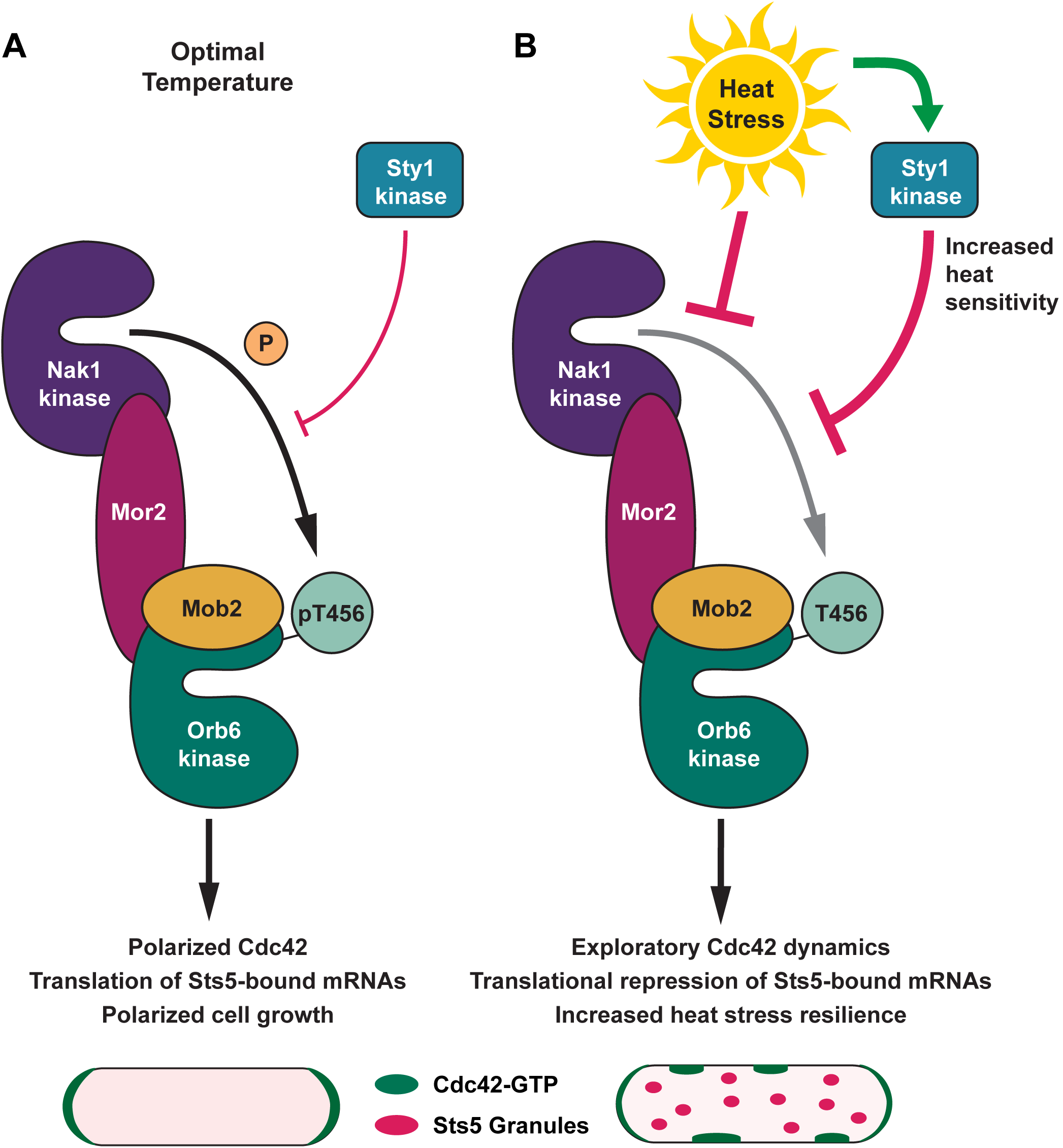
Regulation of NDR kinase Orb6 by temperature and MAP kinase Sty1. (A) Orb6 kinase activity at optimal, not stressful temperatures (25°C) promotes the canonical Cdc42 polarity module (via regulation of RNA Binding protein Sts5), and polarized cell growth. (B) Upon heat stress, MAP kinase Sty1 is strongly activated and inhibits Orb6 kinase activity, thereby promoting exploratory Cdc42 dynamics, Sts5 granule assembly, translational repression of Sts5-associated mRNAs, and increased heat stress resilience. The induction of exploratory Cdc42 dynamics also involves direct phosphorylation of Orb6 substrates by Sty1 Kinase (see Discussion). Inhibition of Orb6 kinase activity during heat stress also engages Sty1-independent regulatory mechanisms, likely affecting other components of the MOR (Morphogenesis Orb6 network) pathway. The activity of MAP kinase Sty1 lowers the threshold temperature that triggers Orb6 inhibition, thereby rendering Orb6 more temperature sensitive and promoting thermotolerance.

### Orb6 downregulation fosters the emergence of exploratory Cdc42 dynamics during heat stress

Cdc42 GTPase is a key regulator of cell polarization and is a highly conserved Rho-family GTPase that regulates cell polarity, actin cytoskeleton organization, and vesicle trafficking. In mammalian myoblasts, heat stress reduces cell migration, at least in part through *CDC42* downregulation (Lu *et al*., 2023). In fission yeast, under normal conditions, active Cdc42-GTP oscillates between cell tips to promote polarized growth (Das *et al*., 2012). Upon environmental stress (including heat), Cdc42 adopts an exploratory activation pattern along the cell membrane, deviating from its usual tip-localized oscillations (Vjestica *et al*., 2013; Chen *et al*., 2019; Salat-Canela *et al*., 2021; Haupt, Ershov and Minc, 2018). Here, we show that during both mild and severe heat stress, Orb6 kinase is inhibited. We previously showed that Orb6 inhibition leads to dephosphorylation of Cdc42 regulators Gef1 and Rga3, and their release from the grip of 14-3-3-protein Rad24 (Das *et al*., 2015; Doyle *et al*., 2025), thereby switching on the stress-activated Cdc42 module, which Gef1 and Rga3 are part of (Salat-Canela *et al*., 2021; Doyle *et al*., 2025). Our findings provide a mechanistic explanation of the process of exploratory Cdc42 dynamics activation during heat stress. The functional role of exploratory Cdc42 dynamics activation during heat stress, or other stresses, however, is still unclear. Induction of exploratory Cdc42 dynamics during nitrogen deprivation (Chen *et al*., 2019) may have the function to facilitate the search for mating partners (Bendezú and Martin, 2013). It is possible that the onset of Cdc42 exploratory dynamics in other stresses is a byproduct of engaging the same MAP Kinase-Orb6 kinase regulatory axis (Doyle *et al*., 2025). Alternatively, temporary loss of cell polarity could signal a physiological transition for the cell, providing the opportunity to activate and inactivate alternative polarity sites upon recovery from stress. In fission yeast, recovery from a disrupting event, such as actin depolymerization by Latrunculin A exposure, can alter the pattern of Cdc42 distribution and ensuing cell polarity (Mutavchiev, Leda and Sawin, 2016). Finally, disrupting cell polarization, in combination with translational repression (see below) may be part of a general physiological strategy to slow cell growth, in an effort to preserve energy (Salat-Canela *et al*., 2023) and extend cell lifespan (Chen *et al*., 2019).

### Sts5 assembles into RNP granules and promotes thermotolerance

Upon exposure to a variety of external stressors, cells form stress granules that help promote stress resilience and survival. The RNA binding protein Sts5 is highly conserved in eukaryotes and is related to human Dis3L2, which is associated with Perlman’s syndrome and Wilm’s tumor in humans. In fungi, Sts5 is closely related to *Saccharomyces cerevisiae* and *Candida albicans* Ssd1, and *Neurospora crassa* GUL-1 (Astuti *et al*., 2012; Lv *et al*., 2015; Robinson *et al*., 2015; Malecki *et al*., 2013; Toda *et al*., 1996; Vaggi *et al*., 2012; Jansen *et al*., 2009; Kurischko *et al*., 2011). These RNA-binding regulators fine-tune translation of cell wall and morphogenesis-related genes, crucial for stress adaptation, survival, and pathogenicity. In the yeasts *Schizosaccharomyces pombe* and *Saccharomyces cerevisiae*, Sts5 and Ssd1, respectively, bind to 5′ UTRs of mRNAs encoding cell wall proteins, repressing their translation to control cell wall biogenesis (Nuñez *et al*., 2016; Hogan *et al*., 2008). Fission yeast Sts5 also alters the translational landscape of mRNAs involved in polarized cell growth and Cdc42 activity, regulating the Ras1-Scd1-Cdc42 regulatory axis (Chen *et al*., 2019; Nuñez *et al*., 2016). By promoting translational repression during nutritional stress, both Sts5 and Ssd1 influence entry into and recovery from quiescence (Chen *et al*., 2019; Miles *et al*., 2019). Finally, both Sts5 (this paper) and Ssd1 (Mir, Fiedler and Cashikar, 2009) are essential for tolerance to heat. Thus, the conservation of structure and cellular function through such enormous evolutionary distance highlights the crucial role of these RNA binding proteins in cell adaptation and survival.

How are these effects exerted in the cell? These proteins all share the property to undergo liquid-liquid phase separation upon exposure to environmental stress, in a manner that engages their N-terminal Intrinsically Disordered Domains (IDRs). The N-terminal domain is crucial to mediate liquid-liquid phase separation of Sts5, Ssd1 and GUL-1 into RNP granules, and is phosphorylated by conserved NDR kinase Orb6 in *Schizosaccharomyces pombe* (Chen *et al*., 2019), Cbk1 in *Saccharomyces cerevisia*e (Kurischko *et al*., 2011; Jansen *et al*., 2009), and COT-1 in *Neurospora crassa* (Stein et al., 2020) and other kinases (such as *S. pombe* Pak1 (Magliozzi and Moseley, 2021)). We previously showed that serine 86 in the N terminus of Sts5 has a crucial role in regulating Sts5 assembly into RNP granules, by mediating N-terminal Sts5 interaction with 14-3-3 protein Rad24 (Chen *et al*., 2019). Association of Sts5 with Rad24 inhibits Sts5 phase separation into granules and maintains active protein translation of Sts5-associated RNAs (Nuñez *et al*., 2016). Conversely, mutation of serine 86 to alanine (*sts5-S86A*), which prevents phosphorylation at that site, inhibits Rad24 association with Sts5 and thereby promotes RNP granule assembly and decreased translation of Sts5-assicated RNAs (Chen *et al*., 2019). We found that by subtly promoting RNP assembly, the *sts5-S86A* mutation seemingly enhances cell resilience to stress exposure, extending chronological lifespan during nutritional stress (Chen *et al*., 2019). Here, we show that the *sts5-S86A* mutation increases cellular resilience to heat stress, while loss of Sts5 increases heat stress vulnerability. We previously showed that Sts5 alters the translational landscape of the cell by decreasing specific mRNA availability for translation, such as *ssp1* mRNA and *efc25* mRNA ((Nuñez *et al*., 2016); Fig. 2C). Consistent with a role for Sts5 in translational repression during heat exposure, Ssp1 protein levels drastically increase upon heat stress, when *sts5* is deleted (Nuñez *et al*., 2016).

Processing bodies and stress granules are types of cytoplasmic ribonucleoprotein (RNP) granules involved in mRNA regulation. A defining function of P-bodies and stress granules is their dynamism, as these granules quickly assemble upon stress exposure and disperse when stress conditions ameliorate. The difference between P-bodies (processing bodies) and stress granules lies in their composition, formation triggers, and cellular roles. P-bodies are involved in mRNA decay, storage, and translational repression and contain enzymes for mRNA degradation (e.g., decapping enzymes Dcp1/Dcp2) (Sakuno et al., 2004; Ingelfinger et al., 2002; van Dijk et al., 2002; Wang et al., 2002). P-bodies are present under normal conditions but increase in size and number during cellular stress. Stress granules, conversely, form in response to cellular stress (e.g., heat shock, oxidative stress) and are triggered by translation initiation inhibition. Stress granules store untranslated mRNAs to prevent their degradation and help cells recover by allowing mRNAs to re-enter translation once stress resolves (Marcelo *et al*., 2021; Panas, Ivanov and Anderson, 2016).

While it is known that P-bodies and stress granules initially assemble through a process known as liquid-liquid phase separation (Aguzzi and Altmeyer, 2016), much remains to be elucidated regarding the mechanisms and factors involved in their formation. Coalescence of RNA binding proteins into puncta is thought to promote nucleation of both P-Bodies and stress granules (Kroschwald et al., 2015; Lin et al., 2015; Elbaum-Garfinkle et al., 2015; Patel et al., 2015; Kato et al., 2012; Hyman, Weber and Julicher, 2014; Brangwynne et al., 2009; Lee et al., 2013; Brangwynne, 2013; Becker and Gitler, 2015).

During nitrogen or glucose deprivation, Sts5 proteins assemble into puncta that partially co-localize with the P-Body marker Dcp1-mCherry (Chen *et al*., 2019; Nuñez *et al*., 2016). In this paper, we find that also upon mild heat stress (36°C), Sts5 coalesces into granules that colocalizes with P-body marker Dcp1. At the heat stress temperature 36°C, we do not observe formation of detectable Pabp1-containing stress granules. Upon severe heat stress (42°C) when visible stress granules form, Sts5 also co-localize with stress granules marker, Pabp1. Consistent with the fact that both P-Bodies and RNA binding proteins are thought to promote the assembly of stress granules (Buchan, Muhlrad and Parker, 2008; Wheeler et al., 2016), we find that loss of *sts5* strongly reduces the number of stress granules and decreases cell survival during heat shock. Conversely, Orb6 inhibition prior to heat exposure, which promotes Sts5 coalescence into granules, (Nunez *et al*., 2016; Chen *et al*., 2019), substantially increases the amount of stress granules formed in the cell during severe heat stress (42°C) and drastically increases resilience to heat stress and heat shock. However, since increase in thermo-resilience is much greater for Orb6 inhibition than for the s*ts5-S86A* mutation, it is likely that Orb6 has additional roles in thermotolerance which are independent of Sts5.

Furthermore, since Orb6 kinase inhibition alone is not sufficient to drive stress granule formation, it is likely that additional factors and regulatory controls play a role in driving Sts5-mediated stress granule formation. For example, other factors such as the vigilin homologue Vgl1 (Wen *et al*., 2010), signaling factors such as calcineurin (Higa *et al*., 2015), and protein kinase C Pck2 (Kanda *et al*., 2021) also associate with stress granules in response to thermal stress, and stress granule formation is also stimulated in response to heat stress by the RNA-binding protein Nrd1 (Satoh *et al*., 2012).

### The MAP kinase Sty1-NDR kinase Orb6 regulatory axis regulates Cdc42 dynamics and RNP granule assembly during heat stress

Conserved MAP kinase Sty1 is the stress-activated MAP kinase (SAPK) in fission yeast. It is the central component of the Wis1–Sty1 MAPK pathway, analogous to mammalian p38/JNK pathways. Heat stress transiently activates Sty1 through its dual phosphorylation by Wis1 at two conserved residues, Threonine 171 and Tyrosine 173, and leads to Sty1 nuclear accumulation (Degols, Shiozaki and Russell, 1996; Smith *et al*., 2002). Sty1 phosphorylation during heat stress is regulated by threonine and tyrosine phosphatases, where tyrosine phosphatases, Pyp1 and Pyp2, are inhibited by heat contributing to Sty1 activation and threonine phosphatases, Ptc1 and Ptc3, work to attenuate Sty1 activity (Nguyen and Shiozaki, 1999). *Sty1Δ* mutants (Sty1 deletion) show severe thermo-sensitivity, losing viability rapidly compared to wild type (Yamada *et al*., 1997). Sty1 activation is essential for survival under both severe heat stress conditions (≥42°C) (Berlanga *et al*., 2010; Yamada *et al*., 1997; Degols, Shiozaki and Russell, 1996), as well as mild heat stress temperature 37°C (Yamada *et al*., 1997).

Our results indicate that Orb6 activity decreases upon exposure to heat stress and heat shock. We recently reported that MAP kinase Sty1 negatively regulates Orb6 during nitrogen starvation (Doyle *et al*., 2025) providing a mechanistic understanding as to why Sty1 activation elicits exploratory Cdc42 dynamics, a behavior that may facilitate ensuing cell mating (Bendezú and Martin, 2013; Merlini *et al*., 2016). We showed that Sty1 inhibits Orb6 kinase, thereby releasing the Orb6 substrates Cdc42 GEF Gef1 and Cdc42 GAP Rga3 from their association with 14-3-3-protein Rad24 (Doyle *et al*., 2025), Sty1 also directly phosphorylates Gef1 and Rga3, likely activating their enzymatic activity (Salat-Canela *et al*., 2021). This “coherent feedforward” mechanism promotes exploratory Cdc42 dynamics during stress response (Salat-Canela *et al*., 2021; Doyle *et al*., 2025).

In this paper, we show that this Sty1-NDR kinase Orb6 regulatory axis has a function in other stress responses, namely heat stress. Further, our data indicate that, through Orb6, Sty1 also modulates the phase separation properties of mRNA binding protein Sts5, and therefore the coalescence of P-bodies and stress granules in response to heat exposure.

Interestingly, our observations show that loss of Sty1 completely suppresses the effects of heat dependent Orb6 inhibition (the induction of Cdc42 exploratory dynamics, Gef1 dephosphorylation, Sts5 aggregation) during moderate heat stress at 36 °C, but not under heat shock conditions at 42°C. Experimentally, these findings suggest a “temperature-specific” control of Orb6 kinase by Sty1 kinase, namely that the Sty1 function in Orb6 inhibition is only effective at sublethal 36 °C temperatures, but not at 42°C heat stress temperatures.

### The role of Orb6 T456 c-terminal hydrophobic motive in thermal sensation

To explore the mechanism behind this effect, we tested the role of the C-terminal hydrophobic motif of Orb6 kinase in thermal sensation. The highly conserved hydrophobic motif (HM) is a critical regulatory element in the AGC family of protein kinases. It contains a phosphorylatable serine or threonine residue (in Orb6, Threonine 456) flanked by hydrophobic amino acids. Phosphorylation of the HM is essential for stabilizing the active conformation of the kinase, facilitating proper alignment of catalytic residues for efficient substrate phosphorylation, and strongly correlates with kinase activity (Millward, Hess and Hemmings, 1999; Hergovich *et al*., 2006; Hergovich, 2013; Jansen *et al*., 2006; Brace, Hsu and Weiss, 2011; Parker *et al*., 2020).

We discovered that the level of HM T456 phosphorylation in Orb6 kinase is temperature-dependent, in a manner that is inversely correlated with increasing temperatures. We find that upon loss of Sty1, the overall levels of Orb6-T456 phosphorylation substantially increase, an effect that is particularly striking at 25°C and 36°C, indicating that Sty1 has a role in the negative control of Orb6-T456 phosphorylation. Further, we find that Orb6-T456 phosphorylation decreases during mild heat stress and severe heat stress in both control or *sty1Δ* strains, suggesting that a Sty1-independent mechanism also mediates Orb6 HM dephosphorylation during heat stress and heat shock. The combined effect is that in *sty1Δ* cells exposed to heat stress at 36°C, the level of Orb6 T456 phosphorylation is comparable to wild-type control cells grown in unstressed conditions at 25°C. Consistent with the idea that the activity of Orb6 remains high in *sty1Δ* cells at 36°C, Gef1 phosphorylation, Cdc42 distribution and Sts5 localization are unperturbed, and cells appear unresponsive to heat stress. Overall, these results indicate that Orb6 kinase inhibition by Sty1 kinase increases heat sensitivity during mild heat stress (36°C), promoting assembly of Sts5 into granules (Fig.1E, 4A), coalescence of Sts5 granules with P-bodies (Fig. S1), and downregulation of Sts5-associated mRNA translation (Nuñez *et al*., 2016). Thus, this finely tuned regulatory network ensures the inhibition of Orb6 during mild heat stress (36°C), when stress granules do not form, but when reprogramming of the translational landscape, and morphogenetic changes at the membrane may be necessary for survival.

It is unclear what could be the Sty1-independent mechanism of temperature-sensing that leads to Orb6 inactivation. One possible mechanism is that Nak1, the kinase upstream of Orb6, is directly inhibited by heat stress. Another possible mechanism is that heat affects protein structure of the Orb6 regulatory complex, perhaps the association of the co-activator Mob2 to Orb6, or the interaction of the Mor2 scaffold with Nak1 and Orb6. Direct inhibition of the Mob2-Orb6 complex by an inhibitory protein is also possible, as detailed for the *S. cerevisiae* protein Lre1, which does not however have homologues in *S. pombe* or higher eukaryotes (Mancini Lombardi *et al*., 2013; Versele and Thevelein, 2001). Future experiments will address these possibilities.

In conclusion, our study highlights the role of the MAP kinase Sty1-NDR kinase Orb6 regulatory axis in promoting thermotolerance during environmental heat exposure. Thermotolerance refers to the ability of an organism, tissue, or cell to withstand elevated temperatures without sustaining irreversible damage. The acquisition of thermotolerance has enormous implications for the survival of crops, the spread of pathogenic organisms to different latitudes, or the success of disease treatments, such as during cancer hyperthermia therapy to enhance chemotherapy or radiation sensitivity. Thus, these findings may have implications in higher eukaryotes due to the highly conserved nature of NDR kinases across evolution.

## Materials and Methods

### Strains and Growth Medium

*Schizosaccharomyces pombe* strains used in this study were derived from wild-type strains 972 or 975 and are displayed in Table S1. *S. pombe* was maintained in yeast extract plus supplements (YES) or Edinburgh minimal medium supplemented with 0.5% ammonium chloride (EMMN) and additional supplements as needed (Histidine, Leucine, Adenine, or Uracil) at a concentration of 225 mg/L. All strains were cultured at 25°C unless otherwise noted. Liquid cultures were incubated at 180 RPM in a shaking incubator, and all cultures were diluted daily such that they were maintained in logarithmic growth for a minimum of 8 generations prior to performing experiments.

### Fluorescence Microscopy

For experiments utilizing microscopy, samples were imaged with an Olympus BX61 fluorescent microscope using appropriate filters. Exposure times range from 1000-2500ms for each experiment and remain consistent among all samples for that experiment. Images were acquired and processed using Intelligent Imaging Innovations SlideBook image analysis software (Version 6.0.4; Denver, CO) and prepared with Fiji (Schindelin *et al*., 2012).

### RNP Granule Induction

To determine localization of Sts5 puncta relative to stress granules, we introduced the marker Pabp1-DsRed, constructing a *sts5-3xGFP pabp1-DsRed* strain (FV2361). We tested the localization of Sts5 relative to stress granules upon heat stress at 36°C or 42°C for 30 minutes. To test the effects of glucose limitation, cells were grown in minimal medium (EMM) containing 2% glucose for at least 8 generations, and then were centrifuged, washed with the appropriate medium, and resuspended either in EMM lacking glucose or in EMM with 2% glucose. Images are representative of samples from three independent experiments.

### RNA Fluorescent In Situ Hybridization (RNA FISH)

Localization of *ssp1* mRNA was determined using RNA-FISH as previously described, with modifications according to Nunez *et al*. (Nunez *et al*., 2016; Nilsson and Sunnerhagen, 2011; Heinrich *et al*., 2013; Brengues and Parker, 2007). *sts5-3xGFP pabp1-DsRed* cells (FV2361) were cultured in EMM lacking glucose for 20 minutes prior to fixation and hybridization of RNA was performed with 20-mer DNA oligonucleotides (Stellaris) labeled with Quasar 705 fluorochromes.

### Glucose Starvation and Orb6 Inhibition Assays

To determine if Sts5 was required for the formation of stress granules, a glucose starvation assay was performed with *sts5^+^ pabp1-DsRed* (FV1684) and *sts5Δ pabp1-DsRed* (FV3192) mutants. Samples were cultured to log phase in minimal medium (EMM) and were washed once with 5 mL minimal medium lacking glucose. Cells were then resuspended in 5 mL minimal medium lacking glucose and were incubated at 25°C and 180 RPM in a shaking incubator for 20 minutes. Following incubation, 1 mL aliquots were centrifuged at 4000 RPM for 1 minute and resuspended in a small amount of residual medium. Microscopy was performed as previously described and the number of stress granules was manually counted. This experiment was performed in triplicate, and n=90 cells per strain per strain per condition in total.

For Orb6 inhibition assays, *orb6-as2 pabp1-DsRed* (FV3226) and *pabp1-DsRed* control (FV1684) strains were cultured to log phase in minimal medium containing adenine. 1-NA-PP1 was added to samples at a final concentration of 50 µM. For assays in which Orb6 was inhibited and then cells were heat shocked, 1-NA-PP1 was added to cells 15 minutes prior to heat shock, whereas DMSO was added for control cells. Samples were then shifted to a 42°C shaking water bath incubator for 20 minutes. Additional controls were maintained at 25 °C. Microscopy was performed as previously described and stress granules were manually counted. This experiment was performed in triplicate, and n = 90 cells per strain per condition in total.

### Heat Shock Survival Assay

*Wild-type* (FV2644), *sts5Δ* (FV2674), *sts5-HA* (FV2645), and *sts5-S86A-HA* (FV2649) strains were cultured in EMM and maintained in log-phase for a minimum of eight generations. Each strain was diluted to O.D._595nm_ = 0.2 (approximately 4×10^6^ cells/mL) in 5 mL EMM and allowed to grow at this density for one hour at 25°C shaking at 180 RPM. Samples were either maintained at 25°C or heat shocked at 48°C for 15 minutes. Following heat shock, experimental and control samples were each serially diluted, plated in triplicate on EMM plates, and incubated at 25°C for approximately 72 hours. Colony forming units per mL (CFU’s/mL) were calculated for each condition, and percent survival was calculated by dividing CFU’s/mL for heat shocked replicates by their respective untreated control CFU’s/mL. Percent survival of each technical replicate was normalized to the average untreated control percent survival. The average percent survival for each strain is as follows: *wt*, 1.8%; *sts5Δ*, 0.2%; *sts5-HA*, 1.3%; and *sts5-S86A-HA*, 4.0%. This experiment was performed in biological triplicate, and data was normalized to the average wild-type percent survival for each independent experiment and subsequently graphed as the relative survival rate. To determine the impact of intermediate to long-term restrictive temperature on these strains, this experiment was repeated with modifications. In this case, samples were incubated at 25°C or 36.5°C for 6 hours. Following incubation, cells were serially diluted and plated in triplicate, and incubated at 25°C for approximately 72 hours. Calculation of CFU’s/mL and normalization of data was performed as previously described, and this experiment was performed in biological triplicate. The average percent survival for each strain (normalized to the control at 25°C) is as follows: *wt*, 125.6%; *sts5Δ*, 65.2%; *sts5-HA*, 120.8%; and *sts5-S86A-HA*, 179.9%.

For heat shock survival analysis of the *orb6-as2* strain (FV2527) and its respective wild-type control (FV2530), cells were initially pre-cultured in minimal medium supplemented with adenine at 180 RPM and 32°C. In order to repress the expression of *orb6-as2* the culture was split into two sets, either with or without thiamine at a final concentration of 15 µM and was allowed to grow for ∼16 hours. Cells were diluted to an equivalent O.D._595nm_ of 0.2 (4×10^6^ cells/mL) in appropriate media, and cells were allowed to grow for an additional 2 hours. Samples were then heat shocked in a 48°C shaking water bath for 15 minutes while controls were maintained at 32°C. All samples were then serially diluted and plated on minimal medium supplemented with adenine, but lacking thiamine, for subsequent quantification of CFU’s/mL as previously described. The average percent survival for each strain is as follows: *wt* - Thiamine, 0.1%; *wt* +Thiamine, 0.1%; *orb6-as2* - Thiamine, 0.1%; and *orb6-as2* +Thiamine, 18.9%.

### Temperature Dependent Analysis of Sts5 Puncta

*sts5-3xYFP* (FV2518) and *sts5-S86A-3xYFP* (FV2522) strains were used to determine whether Sts5 puncta are induced at lower temperatures when Sts5 is hyperactive as compared to a wild-type allele. Each strain was cultured to log-phase in EMM, diluted to O.D._595nm_= 0.1, and cells were allowed to grow for a minimum of 1 hour at 25°C. Aliquots of each strain were created, and samples were heat shocked at 29°C, 32°C, 35.5°C, or 42°C for 30 minutes. Control cells were maintained at 25°C throughout the experiment. Following exposure to heat shock, samples were imaged using an Olympus BX61 microscope. To determine differences in number of puncta, 20 cells per strain were manually scored for Sts5 (YFP) puncta at each temperature.

### Development of anti-pOrb6-T456 phospho-specific antibody

The anti-pOrb6-T456 phospho-specific antibody was custom-made by 21st Century Biochemicals against a synthetic peptide corresponding to the phosphorylated Thr-456 (amino acids 449-462, NLAFLGYT*YKKFNY) of the fission yeast Orb6 protein. To detect pOrb6-T456, we used a strain expressing N-terminally tagged HA-Orb6 *(HA-orb6-as2)*, integrated in single copy behind the *nmt1* promoter, since increased expression of Orb6 allows for better detection of pOrb6-T456, as compared to C-terminally tagged Orb6-HA or endogenous Orb6 where the detection signal is too low.

### Protein extraction and western blot analysis

Protein extraction was modified from previously described protocol(Matsuo *et al*., 2006) and performed as previously described (Doyle *et al*., 2025). ∼7.5 × 10^7^ total cells growing exponentially were harvested and cell pellets were resuspended in 300μL of H_2_O with protease and phosphatase inhibitor cocktail (Halt™ Protease and Phosphatase Inhibitor Cocktail (100X)) and transferred to 1.5mL microcentrifuge tube. 300μL of 0.6M NaOH with inhibitor cocktail (see above) was added to the cells and resuspended. Cells were left to incubate for 5 minutes at room temperature, inverting the tubes 2-3 times halfway through incubation. Cells were then centrifuged at 2300 RCF for 2 minutes. The supernatant was discarded, and the pellet was resuspended in 75μL modified SDS buffer (60mM Tris-HCl [pH 6.8], 4% 2-Mercaptoethanol, 4% SDS, 5% glycerol) with inhibitor cocktail (see above) and boiled at 98°C for 3 minutes. Cells were placed on ice and centrifuged at 4°C at 3500 RCF for 1 minute. 65μL of the supernatant (protein extract) was collected with 60μL stored at −80°C immediately, and 5μL used for downstream protein quantification assay. To quantify protein, the RC DC protein assay kit was used as it is compatible with the concentration of reducing agents and detergents present in the modified SDS buffer.

Standard western blotting procedures were performed as follows: protein was separated using SDS-PAGE and transferred onto a nitrocellulose membrane. The membranes were probed with antibodies of interest and visualized through fluorescent detection on Li-Cor Odyssey CLx. Gef1-3YFP strains were used in detecting total Gef1 protein with anti-GFP (Roche, CAT#: 11814460001; RRID:AB_390913; dilution factor (DF): 1:1000) and phosphorylated Gef1-S112 was detected using a custom-made previously described antibody(Chen *et al*., 2019). HA-Orb6as2 strains were used in detecting total Orb6 protein with anti-HA (Biolegend, CAT#901501; RRID:AB_2565006; DF: 1:3000) and phosphorylated Orb6-T456 was detected using a custom-made antibody (see subsection “Development of anti-pOrb6-T456 phosphospecific antibody” in Methods section). Anti-β-Actin (Abcam, CAT#: AB8224; RRID:AB_449644; DF: 1:4000) was used as a loading control for all experiments. Two-color detection was used with using LiCor IRDye secondary antibodies anti-rabbit 800CW (Li-Cor, CAT#: 926-32211; RRID:AB_621843; DF: 1:15000) for anti-pGef1-S112 or anti-pOrb6-T456 and anti-mouse 680RD (Li-Cor, CAT#: 102673-408; RRID:AB_10956588; DF: 1:15000) for anti-GFP (Gef1-3YFP) or anti-HA (HA-Orb6as2). Quantification of the blots was performed using Image Studio software. To quantify, a rectangle was drawn around the signal band of interest and pixel intensity was recorded. To remove background signal, background settings were set to median, segment: top and bottom, border width: 3. With the pixel intensity measurements, a ratio was determined between pGef1-S112 or pOrb6-T456 (fluorescent channel 800) and total Gef1-3YFP or total HA-Orb6as2 (fluorescent channel 700) to measure how much total Gef1-3YFP protein is phosphorylated at S112 or total HA-Orb6as2 protein is phosphorylated at T456. Data was normalized by creating a ratio with all values with the average ratio of control strains and/or conditions. Similar quantification was performed to measure ratio of Atf1 to β-Actin during heat stress in control cells and *sty1* deletion mutants using anti-Atf1 (Abcam, CAT#: AB18123; RRID:AB_444264; DF: 1:2000) and anti-β-Actin (Abcam).

## Quantification and statistical analysis

Data are presented as described in legends. Experiments were completed in independent biological triplicates. A two-tailed unpaired Student’s t test was used to assess statistical significance between two groups. One-way or two-way analysis of variance (ANOVA) followed by appropriate post hoc test was applied to evaluate the difference between more than two groups (one-way) or more than 2 groups and more than one condition (two-way). Statistical analyses and visualization were performed with GraphPad Prism 10 (GraphPad Software, San Diego, CA). p-value < 0.05 was set as the threshold for statistical significance. Power analyses were performed using G*Power to determine minimum sample size(Faul *et al*., 2007; Kang, 2021).

## Data and Resource Availability

Data and resource availability: All relevant data and details of resources can be found within the article and its supplementary information.

