## Supplementary files for "Temperature Specific Regulation of NDR Kinase Orb6 by MAP kinase Sty1 to Promote Heat Stress Resilience"

### Supplementary Figures and Legends

| Strain | Genotype | Source |
|---|---|---|
| FV1684 | *pabp1-DsRed::KanMX6* | (Nilsson and Sunnerhagen, 2011) |
| FV2267 | *sts5-3xGFP::NatMX6 dcp1-mCherry::hph* | (Nuñez *et al.*, 2016) |
| FV2361 | *sts5-3xGFP::NatMX6 pabp1-DsRed::KanMX6* | This Study. |
| FV2518 | *sts5Δ::NatMX6 sts5-HA::KanMX6* | (Chen *et al.*, 2019) |
| FV2522 | *sts5Δ::NatMX6 sts586A-HA::KanMX6* | (Chen *et al.*, 2019) |
| FV2527 | *orb6Δ::ura4+ pJK148orb6-as2::leu1+* | (Chen *et al.*, 2019) |
| FV2530 | *Wild Type, PN972* | (Leupold, 1949) |
| FV2644 | *Wild Type, PN975* | (Leupold, 1949) |
| FV2645 | *sts5Δ::NatMX6 sts5-HA::KanMX6* | (Chen *et al.*, 2019) |
| FV2649 | *sts5Δ::NatMX6 sts5S86A-HA::KanMX6* | (Chen *et al.*, 2019) |
| FV2674 | *sts5Δ::KanMX6* | (Chen *et al.*, 2019) |
| FV3192 | *sts5Δ::NatMX6 pabp1-DsRed::KanMX6* | This Study. |
| FV3226 | *orb6Δ::ura4+ pjk148orb6-as2::leu1+ pabp1-DsRed::KanMX6* | This Study. |
| FV3527 | *CRIB-GFP::ura+ ade+ leu+* | (Chen *et al*, 2019), this study |
| FV3712 | *CRIB-GFP::ura+ sty1Δ::kanMX6 ade+ leu+* | This study |
| FV2206 | *gef1Δ::natMX6 gef1-3YFP::kanMX6 ade+ leu+ ura+* | (Chen *et al*, 2019) |
| FV3634 | *gef1Δ::ura+ gef1-3YFP::kanMX6 sty1Δ::kanMX6 ade+ leu+* | This study |
| FV2444 | *sts5Δ::natMX6 sts5-3YFP::kanMX ade6- leu1-32 ura3-D18* | This study |
| FV3797 | *sty1Δ::kanMX6 sts5Δ::natMX6 sts5-3YFP::kanMX ade6- leu1-32 ura3-D18* | This study |
| FV3343 | *nak1-ts (orb3-167) orb6Δ::ura+ pJK148HA-orb6-as2::leu+ ade+* | This study |
| FV3602 | *orb6Δ::ura+ pJK148HA-orb6-as2::leu+ ade+* | This study |
| FV3642 | *sty1Δ::kanMX6 orb6Δ::ura+ pJK148HA-orb6-as2::leu+ ade+* | This study |

**Supplemental Table 1:** *S. pombe* strains used in this study.

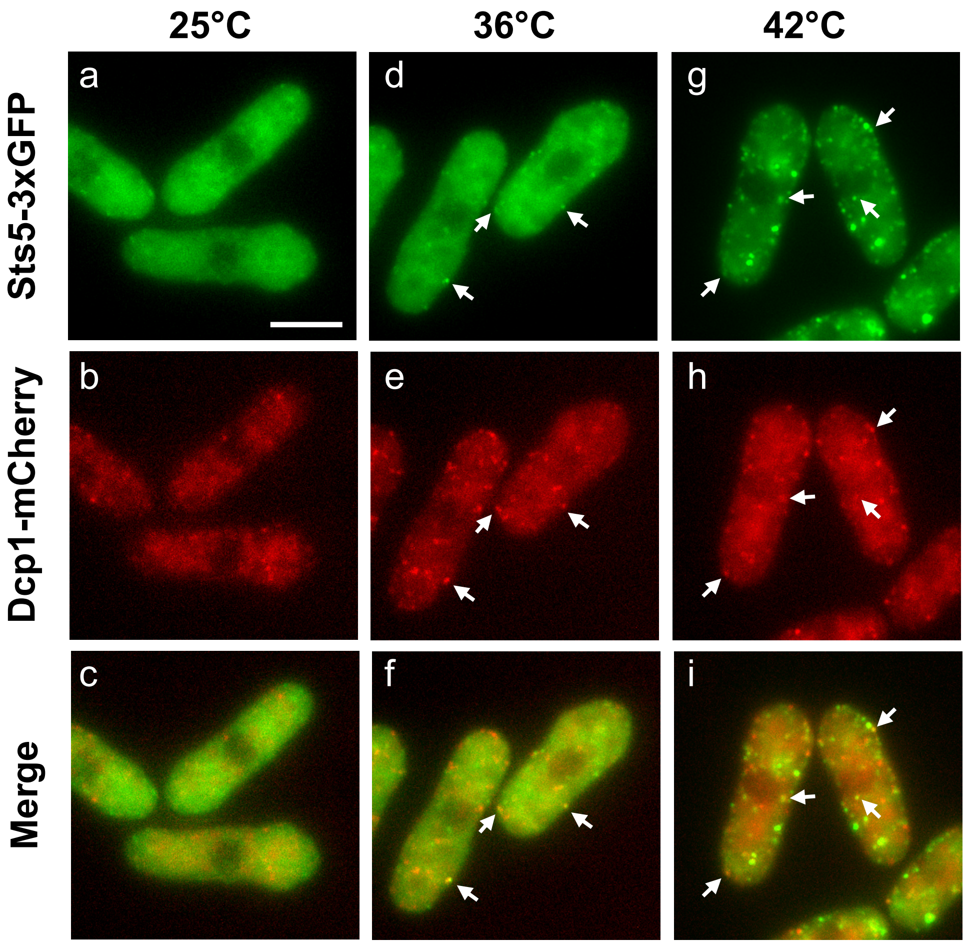


**Supplementary Figure 1: Sts5-3xGFP co-localizes with P-bodies upon heat stress.** Sts5-3xGFP Dcp1-mCherry cells were cultured in YES medium and heat stress at 36ºC or 42ºC for 30 minutes. After heat shock, Sts5-3xGFP formed cytoplasmic puncta which partially co-localized with the P-body marker Dcp1-mCherry at both 36ºC (d-f) and 42ºC (g-i), whereas the cells at control temperature displayed little to no formation of P-Bodies or Sts5 puncta (a-c). Images are deconvolved projections from Z-stacks (6 slices) separated by a step size of 0.3 μM (Bar = 5 μM).

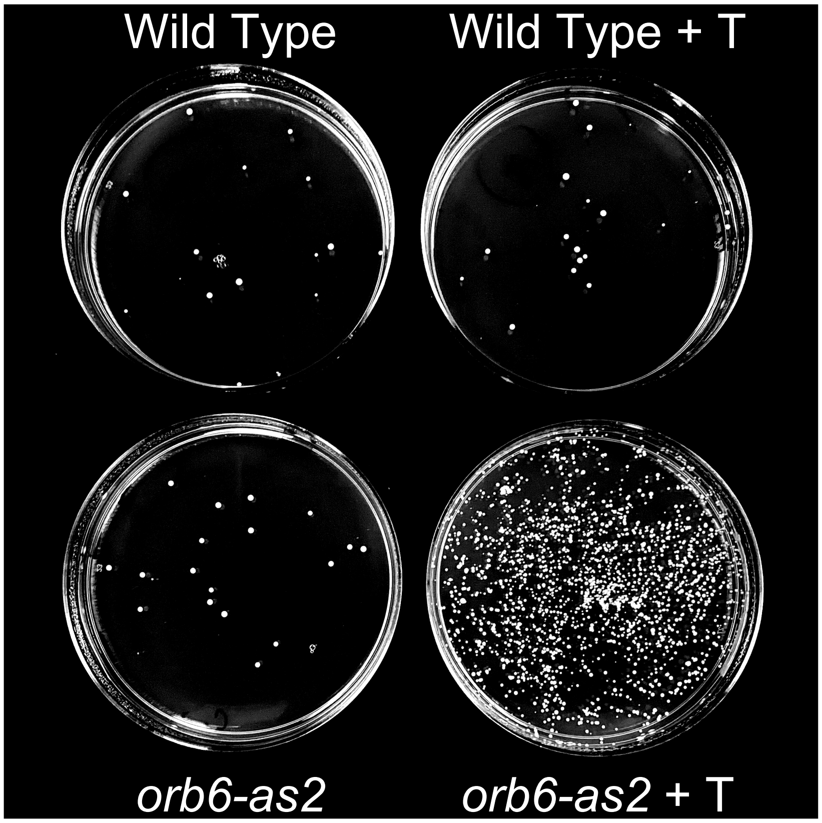


**Supplemental Figure 2: Downregulation of Orb6 promotes survival after heat shock.** Representative image of the orb6-as2 heat shock assay (10-2 dilution) from the heat shocked sample set. Inhibition of Orb6 drastically increases survival (bottom right) after exposure to a 48ºC heat shock for 15 minutes, as compared to untreated or wild-type controls.

**Blot Transparency**

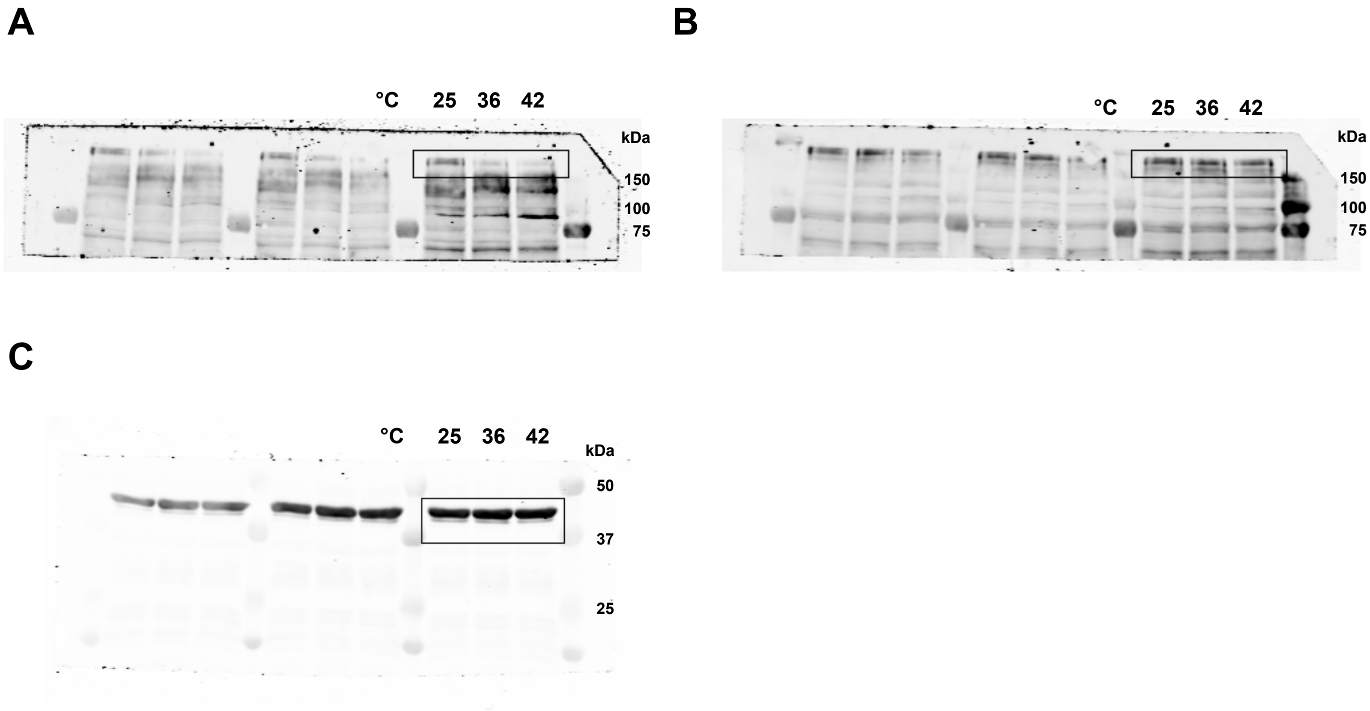


**Uncropped blots from Figure 1C.** (A) anti-pGef1-S112. (B) anti-GFP (total Gef1-3YFP). (C) anti-Actin for A-B.

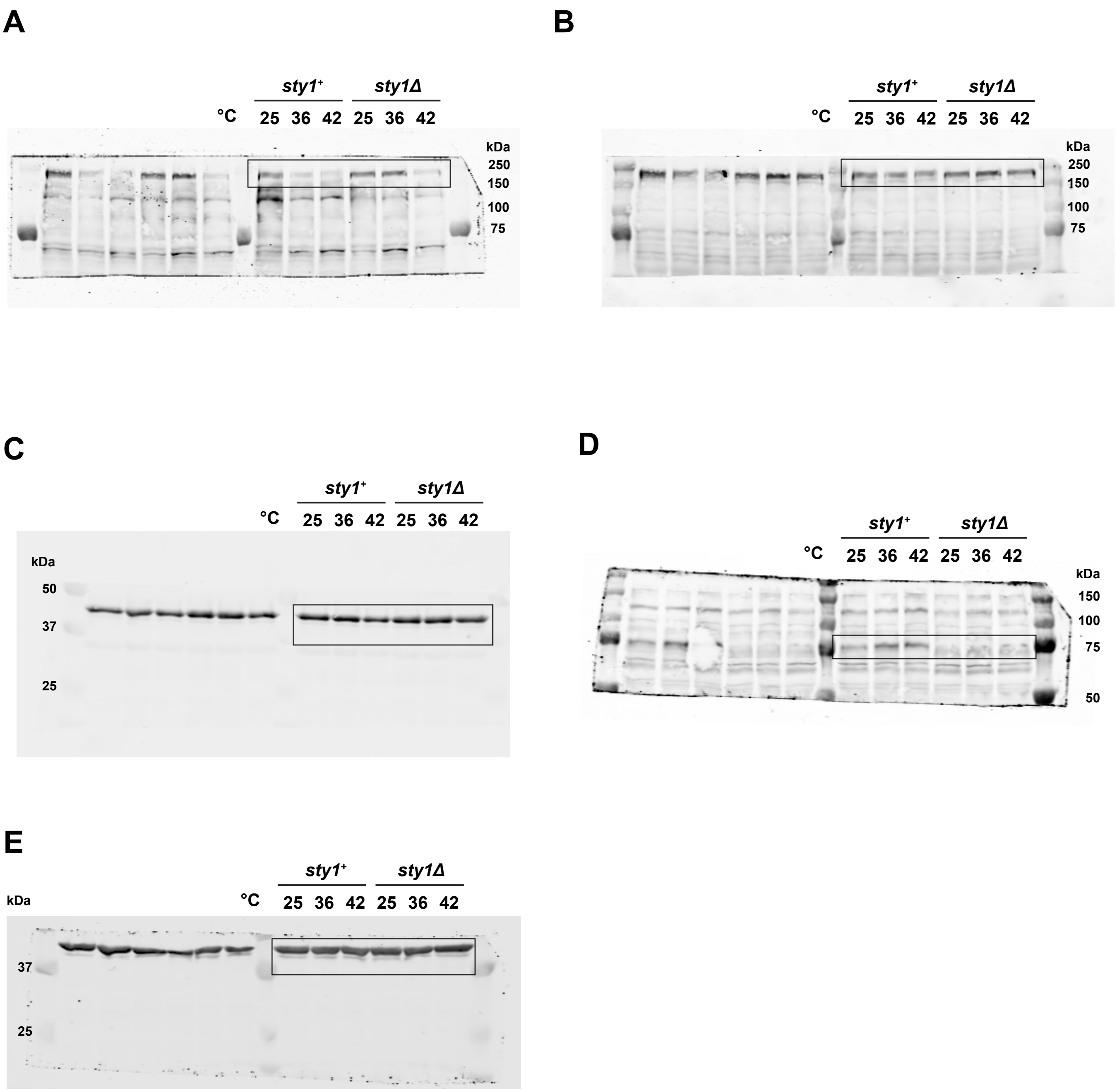


**Uncropped blots from Figure 4G.** (A) anti-pGef1-S112. (B) anti-GFP (total Gef1-3YFP). (C) anti-Actin for A-B. (D) anti-Atf1. (E) anti-Actin for D.

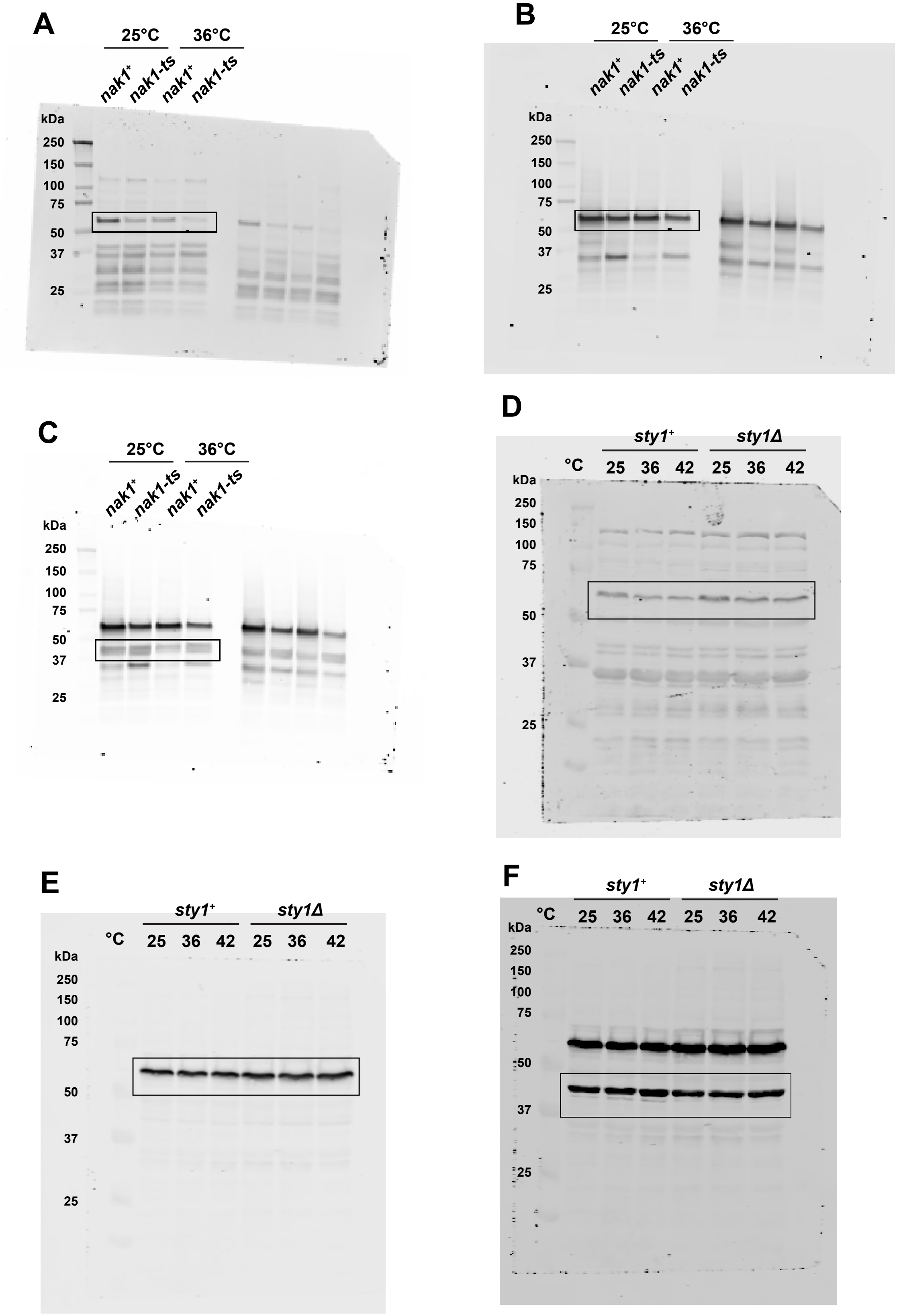


**Uncropped blots from Figure 5.** (A) anti-pOrb6-T456 from Figure 5C. (B) anti-HA (total HA-Orb6as2) from Figure 5C. (C) anti-Actin for A-B. (D) anti-pOrb6-T456 from Figure 5E. (E) anti-HA (total HA-Orb6as2) from Figure 5E. (F) anti-Actin for D-E.
